# Unveiling the Molecular Architecture of T Cells and Immune Synapses with Cryo-Expansion Microscopy

**DOI:** 10.1101/2025.04.15.648816

**Authors:** Florent Lemaître, Olivier Mercey, Isabelle Mean, Elise Paulin, Valérie Dutoit, Jan Rath, Denis Migliorini, Caroline Arber, Paul Guichard, Virginie Hamel, Benita Wolf

## Abstract

Cellular communication is critical for anti-cancer immunity, with tumor cell killing occurring at immunological synapses (IS) formed between effector immune cells and target tumor cells. While optical super-resolution microscopy (SRM) has enlightened the spatial organization of the IS mostly in regular immune cells, visualizing the nanoscale architectural features of IS in its native state, including 3D receptor distribution and the ultrastructural details of the lytic granule release remains challenging. Using cryo-expansion microscopy (cryo-ExM), we unravel the cellular architecture of activated T cells and T cell-target cell pairs. Our approach visualizes actin and microtubule networks during synapse formation, membrane topography, and the distribution of signaling molecules and lytic granules of different types, offering novel insights into IS organization. Finally, we apply U-ExM to glioblastoma tissue, visualizing T cells and their lytic content *in situ*, highlighting its potential for pre-clinical immunotherapy studies.

## Main text

Advanced imaging is crucial for visualizing the complex spatiotemporal regulation of immune signaling^1^, involving tightly regulated interactions between cytotoxic T cells and cancer cells. These interactions form the immunological synapse (IS), a signaling platform that integrates spatial, mechanical, and biochemical signals^2^. During T cell receptor (TCR) signaling, T cells polarize their actin and microtubule cytoskeletons towards the IS, transporting lytic granules that release cytotoxic proteins like Perforin and Granzymes into the target cell^3, 4, 5^. The IS’s spatial organization ranges from large micron-sized actin rings to sub-resolution receptor micro-clusters^6, 7^.

Super-resolution techniques like single molecule localization microscopy (SMLM), structured illumination microscopy (SIM) and stimulated emission depletion (STED) microscopy have been essential to decipher these structures^8, 9, 10, 11^ but these methods often require chemical fixation like paraformaldehyde (PFA) or methanol, which can distort cellular architecture, membrane based organization, and rarely preserve both actin and microtubule cytoskeletons^12, 13, 14^. Live imaging avoids fixation but often lacks resolution for fine structures. Furthermore, while super-resolution microscopy can achieve nanometric resolution, it falls short of capturing the broader cellular context provided by electron microscopy. Consequently, this limits a comprehensive understanding of the IS’s intricate architecture.

To overcome these limitations, we investigated the immune synapse architecture using Cryo-expansion microscopy (Cryo-ExM), which combines cryo-fixation with Ultrastructure-Expansion Microscopy (U-ExM)^15^. Cryo-fixation rapidly freezes cells in a vitreous state and thus preserves their native state, while U-ExM isotropically expands the sample four-fold enabling super-resolution imaging, and combined with pan-labeling N-Hydroxysuccinimide-ester (NHS)^16^ staining, can reveal the cellular context^15^. Cryo-ExM thus provides a robust and straightforward method to visualize the near-native nanoscale organization of human T cells, in resting state and also during their interactions with target cells.

We first set out to test the cytoskeleton preservation ability of cryo-fixation compared to PFA fixation in primary human T cells stained for both actin and tubulin. We found a noticeable improvement in cortical actin structure and microtubule staining upon cryo-fixation, highlighting excellent preservation of these cytoskeletal structures (**Extended Data** Fig. 1a, b). We next applied the expansion procedure using the Cryo-ExM protocol^15^ to human healthy donor primary T cells (**Fig. 1a-c**) and the Jurkat CD4+ T cell line (**Extended Fig. 1c, d**). We first monitored the isotropic expansion of T cells by determining the nuclear cross section value before and after expansion^17^ (**Fig.1d, Extended Data** Fig. 1e ). We confirmed that both Jurkat and primary T cells expanded isotropically as shown previously for other cell types^15, 17, 18, 19^, however, Jurkat cell nuclei expanded slightly less than the gel **(Extended Data** Fig. 1e).

**Figure 1:**
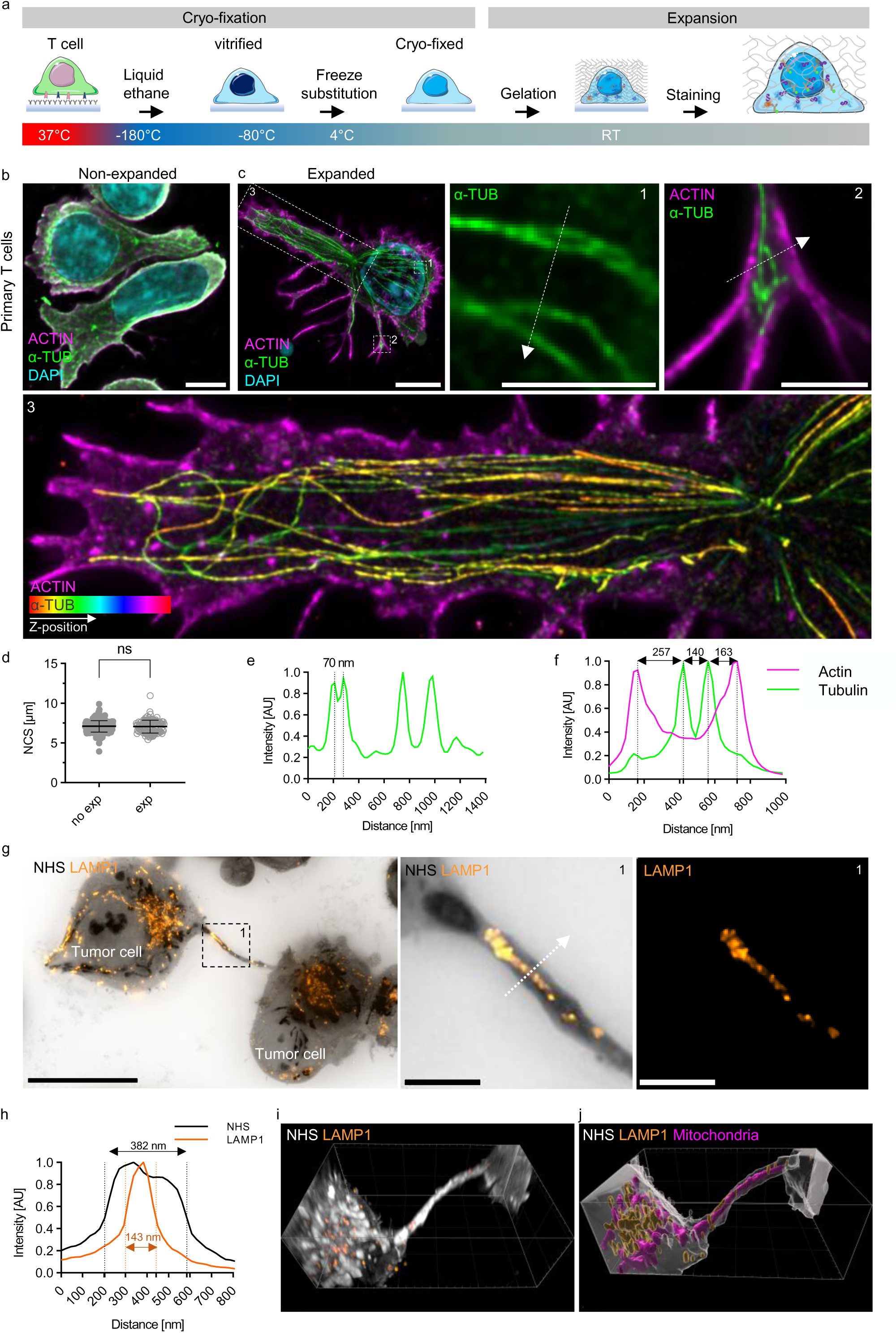
Cryo-ExM preserves the T cell nanometric ultrastructure. **a**, Schematic of the cryo-expansion protocol, adopted from^15^. **b, c**, Representative widefield images of cryo-fixed primary T cells before (non-expanded, **b**) and after (expanded, **c**) expansion, immunofluorescently labelled for actin (magenta), α-tubulin (α-TUB, green) and nucleus (DAPI, cyan), imaged with a widefield microscope, scale bar: 2 µm. Inset (1-3) from (c) show preservation of α-TUB (green, inset 1, 2) and actin (magenta, inset 1) and α-tubulin color-code reflecting the image depths over the z-stack (hyperstack, inset 3). Inset 1-2, scale bar: 500 nm; inset 3, scale bar: 2 µm. **d**, measurement of primary T cell nuclear cross section (NCS) before and after expansion, data corrected for expansion factor (4.3-4.5). Data from n = 3 experiments. Mann-Whitney test, mean ± SD is shown. ns: non-significant. **e**, Plot profile of the tubulin pixel intensity (arbitrary unit) along the white dashed arrow (nm) presented in (**c**) inset 1. xy-resolution of 70 nm. **f**, Plot profile of the normalized actin and α-TUB pixel intensity (arbitrary unit) along the white arrow (nm) presented (**c**) in inset 2. Distances (nm) between indicated peaks are shown. **g**, Widefield image of MM.1S multiple myeloma cells forming a nanotube and immunofluorescently labelled with NHS (N-Hydroxysuccinimide ester, inverted grey), and lysosomal vesicles with LAMP1 (orange), after cryo-ExM, scale: 10 µm. The inset (1) of (**g**), is shown as stained for either NHS and LAMP1 (middle panel) or LAMP1 only (right panel). Maximum intensity z-stack projection (MIP), scale bar: 500 nm. **h**, Plot profile of the normalized NHS and LAMP1 pixel intensity (arbitrary unit) along the white dashed arrow (nm) presented in inset 1 in (g). The xy resolution at 50% fluorescence intensity is indicated. **i, j**, Imaris 3D reconstruction of (g). (i) shows the fluorescent signal. **j,** 3D Surface segmentation of LAMP1 vesicles (orange) mitochondria (purple) and cell surface (grey) performed on the NHS signal.

Then, we assessed the resolutive power of cryo-ExM on T cells and found that it enables the visualization of the spatial distribution of the actin and microtubule cytoskeleton elements (**Fig. 1c, e, f**) with a lateral optical resolution of 70 nm with a widefield fluorescence microscope, similar to other expansion microscopy methods^20, 21^. –Finally, to evaluate the capacity of Cryo-ExM to preserve fragile membrane-based structures and to reveal the cellular context using NHS-ester pan labeling^22^, we turned to analyzing tunneling nanotubes (TNT) often observed in cancer cells. TNT are communication channels between cells, whose diameter varies between 100 nm to 800 nm^23, 24^, that can span long distances and serve to transmit cellular organelles, ions, peptide MHC complexes or any cellular content. Owing to their dimensions and fragility, imaging TNT is often challenging^25^. We found that Cryo-ExM preserved and allowed visualization of TNT between neighboring tumor cells (**Fig. 1g-j**). Our analysis shows that Lysosomal-associated membrane protein 1 (LAMP1)-positive vesicles are well preserved and traverse TNTs, indicating active material exchange between tumor cells. Furthermore, NHS ester staining effectively reveals cellular context, enabling the 3D segmentation of the cell surface, while also highlighting darker organelles that likely correspond to mitochondria (**Fig. 1g-j** and **Extended Data** Fig. 1f-i). This finding is further validated by co-staining with the mitochondrial marker TOM20 (**Extended Data** Fig. 1f-i**, Extended Data Video 1**). Together, these data demonstrate how Cryo-ExM preserves and enhances imaging of T cell actin and microtubule cytoskeleton and very fine tumor cell features.

We next analyzed centrioles, a microtubule-based structures forming the centrosome previously used as a molecular ruler to assess expansion isotropicity, which also plays a crucial role in immune synapse formation^26, 27^ (**Fig. 2a-i**). Upon TCR engagement and PKCθ signaling, the centrosome polarizes towards the synapse^28, 29^, organizing the subcortical microtubule network for lytic granule delivery and Golgi polarization. The centrosome positions close to the cell cortex in the central IS, but whether one centriole docks to the plasma membrane, like cilia formation, remains unclear^27, 30^. The exact role of the centrosome in synaptic killing and signaling remains to be discovered^28, 29^. Using regular U-ExM^20^ protocol, best amenable to visualize centrioles, we observed that the centriole diameters in Jurkat (252 ± 20 nm) and primary T cells (243 ± 32 nm) were comparable to those in U2OS cells (235 ± 15 nm), with only minor variations (**Fig. 2a, b**). However, their lengths were significantly different, measuring 252 ± 91 nm for Jurkat cells and 374 ± 58 nm for primary T cells, in contrast to the canonical length of 429 nm observed in U2OS osteosarcoma cell^31, 32, 33^ (**Fig. 2a, c**). To further validate these findings, we analyzed the three-dimensional architecture of centrioles from Jurkat and primary T cells available from Openorganelle.janelia.org obtained by focused ion beam milling^34^, confirming that human T cells have shorter centrioles compared to other human cells, like U2OS cells (**Extended Data** Fig. 2a, b) or HeLa cells^32^. To explore the centriolar nanostructure and the roots of this difference, we mapped distal centriolar proteins such as C2CD3, known to regulate centriole length^35^, the sub-distal appendage protein CEP170 involved in microtubule anchoring^36^ and the distal appendage protein CEP164 critical for centriole docking to the plasma membrane^37^. Strikingly, we found that Jurkat T cells lack the three distal proteins, while primary T cells contain C2CD3 and CEP164 but miss CEP170 (**Fig. 2 d-i)**. Given the highly mutated genome of Jurkat cells^38^, it could be that the proteins are not expressed. Western blot analysis of cell lysates demonstrates that CEP170 is most likely present in Jurkat T cells but not localized to centrioles (**Extended Data** Fig. 2c), a result consistent with a recent mass spectrometry study reporting a lack of distal appendage proteins in centrioles isolated from Jurkat cells^39^. Taken together this suggests an altered distal centriolar structure in Jurkat T cells and to a lesser extend also in healthy donor T cells. The impact of altered centriolar structure on centrosome-mediated granule release at the immunological synapse remains unclear. Jurkat T cells, which differ in synapse topology from primary human T cells and lack lytic granule delivery, may reflect centriolar structural contributions that warrant further investigation^40, 41^.

**Figure 2:**
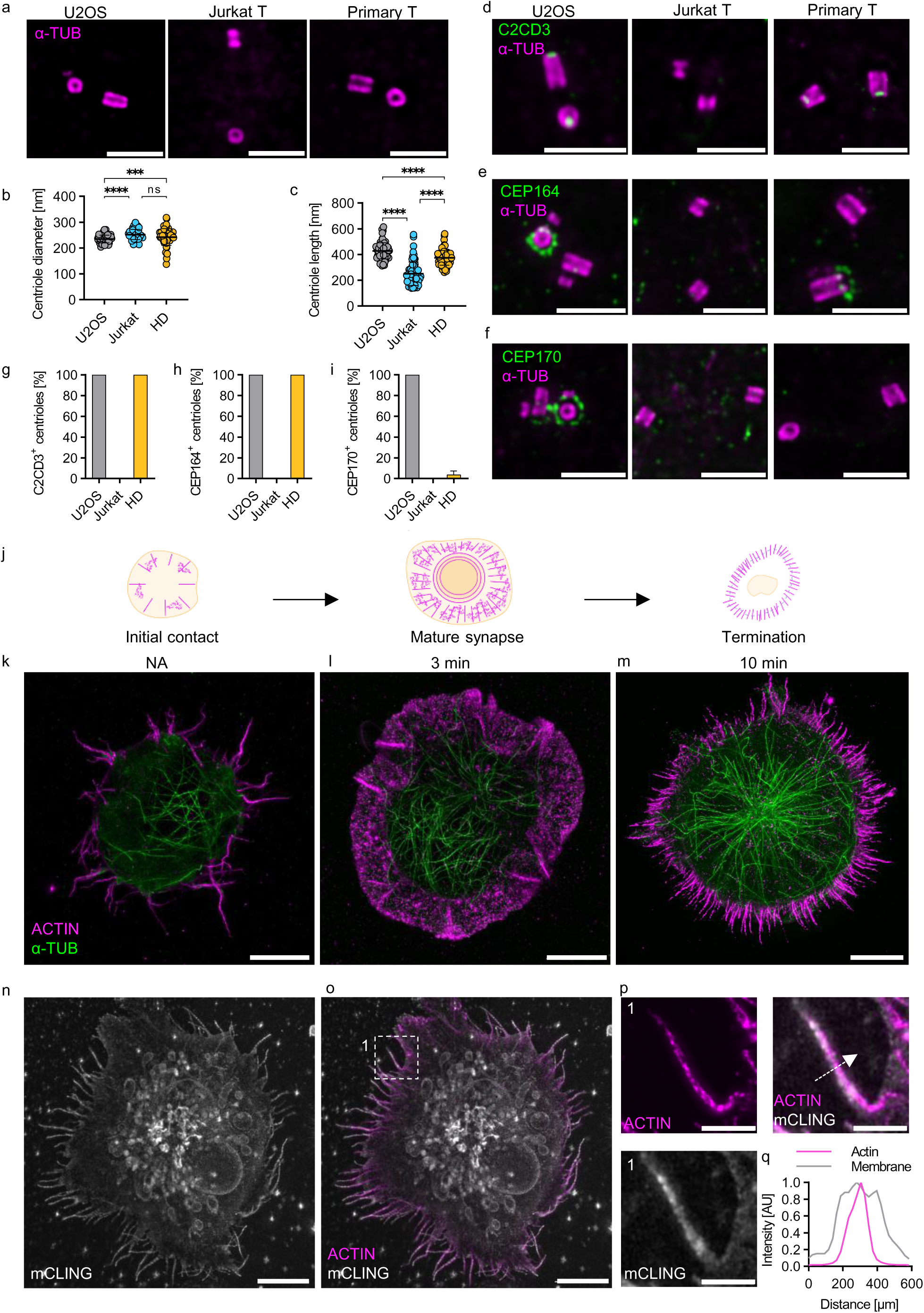
T cell centriolar architecture, IS maturation stages and T cell lipid membranes. All images were taken using a widefield microscope. **a,** U-ExM images of U2OS, activated Jurkat T cell and primary human T cell’s centrioles labeled for α-Tubulin (magenta). Images are MIPs representative from n= 4 independent experiments, scale bar: 1 µm. **b,** centriolar diameter measured from α-tubulin staining for U2OS, Jurkat and primary human T cells. Data are from n = 4 independent experiments, each dot represents one centriole. Kruskal Wallis test with Dunn’s correction, mean ± SD is shown. **c,** centriolar lengths measured from α-tubulin staining for U2OS, Jurkat and primary human T cells. Kruskal Wallis test with Dunn’s correction, mean ± SD is shown. **d-f,** U-ExM images of U2OS, activated Jurkat T cell and primary human T cell’s centrioles labeled for C2CD3 (d), CEP164 (e), CEP170 (f) (in green) and α-Tubulin (magenta). Representative images (MIP) from n = 4 experiments, scale bar: 1 µm. **g-i,** proportion of U2OS, Jurkat and primary human T cells centrioles positive for C2CD3 (g), CEP164 (h) or CEP170 (i) signal from (d-f). Data from n = 4 experiments, mean ± SD is shown. Statistical tests are indicated, ns: non-significant, ***p<0.01, **** p<0.001. **j,** schematic of the immune synapse (IS) maturation stages and actin (magenta) organization over time, adopted from^8^. **k-m,** maximum intensity projections (MIP) of representative Cryo-ExM images of actin (magenta) and α-tubulin (green) organization in Jurkat cells seeded on activating surfaces and illustrating initial contact (k), mature synapse (l) and termination stage (m), scale bar: 5 µm. **n, o,** representative cryo-ExM images (MIP) of an activated Jurkat cell, labelled for mCLING (lipids, grey) (n), or mCLING and Actin (magenta) (o), scale bar: 5 µm **p,** inset 1 in (f) showing the actin cytoskeleton within a membrane labelled microvilli, scale bar: 1 µm.**q**, Plot profile of normalized mCLING (grey) and actin (magenta) pixel intensity (arbitrary unit) along the white arrow (nm) in (**g**).

Next, we focused on the immune synapse (IS). Steps of IS formation encompass initiation, maturation and termination (**Fig. 2j, Extended Data** Fig. 2d-i), with actin and microtubule network rearrangements as landmark features^8^. We set out to investigate whether cryo-ExM could capture these reorganizations in activated Jurkat T cells (**Fig. 2k-m**). Importantly, cryo-ExM allows visualizing the three stages of IS assembly, with utmost preservation of both actin and microtubules, recapitulating the spatial reorganization of these networks upon T cell activation using a widefield microscope. We further explore membrane and actin visualization by co-staining cryo-expanded cells with mCLING^42^, labeling T cell lipid membranes^43^ and actin (**Fig. 2n-p**). We found that membrane structures are preserved, and penetrating actin fibers can be accurately imaged in cell membranes protrusions (**Fig. 2p, q**). Capitalizing on the good preservation of cellular structures using Cryo-ExM, we then monitored the co-stimulatory, adhesion and signaling molecule CD2^44^ and the lymphocyte-specific protein tyrosine kinase (Lck)^45^ (**Fig. 3** and **Extended Data** Fig. 3). CD2 has a dual role as important IS adhesion and signaling molecule^44^ whereas Lck is the major activating kinase of pathways downstream of the TCR^46^. CD2, a small single-pass transmembrane protein of the immunoglobulin superfamily, binds CD58 on target cells. Along with integrin LFA-1, CD2 plays a crucial role in immune synapse formation^2^ and is instrumental in cancer immune evasion when CD58 is downregulated on cancer cells^47^. It localizes to the tip of T cell microvilli^48^ and has been visualized using optical reconstruction from TIRF data and electron microscopy^48, 49^. Here, we show the localization of CD2 on non-activated (**Fig. 3a, Extended Data** Fig. 3a) and activated (**Fig. 3b-d, Extended Data** Fig. 3b) primary CD4 T cells. As expected from the literature, we found CD2 associated with the actin cytoskeleton of microvilli in resting and activated CD4 T cells (**Fig. 3a, b, Extended Data** Fig. 3a, b left panel), and localizing at the periphery of the IS during T cell activation (**Extended Data** Fig. 3b, middle panel), and towards the center of the mature IS (**Fig. 3c, Extended Data** Fig. 3b, right panel). Since cryo-ExM allows 3D volumetric imaging, we could obtain maximum intensity projections of various thicknesses in xy and xz dimensions, allowing for cell side view observation (**Fig. 3d**). As expected, this complete 3D representation of T cells seated on activated surfaces showed an increased spread over time reflected in an enhanced synaptic area and a decreased cell height (**Fig. 3e, 3f**). Finally, analysis of the 3D distribution of CD2 showed increasing amounts at the synapse center over time (**Fig. 3h-m**), with respect to the overall cellular CD2 distribution (**Fig. 3h**) as well as with respect to the actin fluorescence intensity (**Fig. 3j-m**), possibly due to internalization of CD2-containing lipid rafts ^50^.

**Figure 3:**
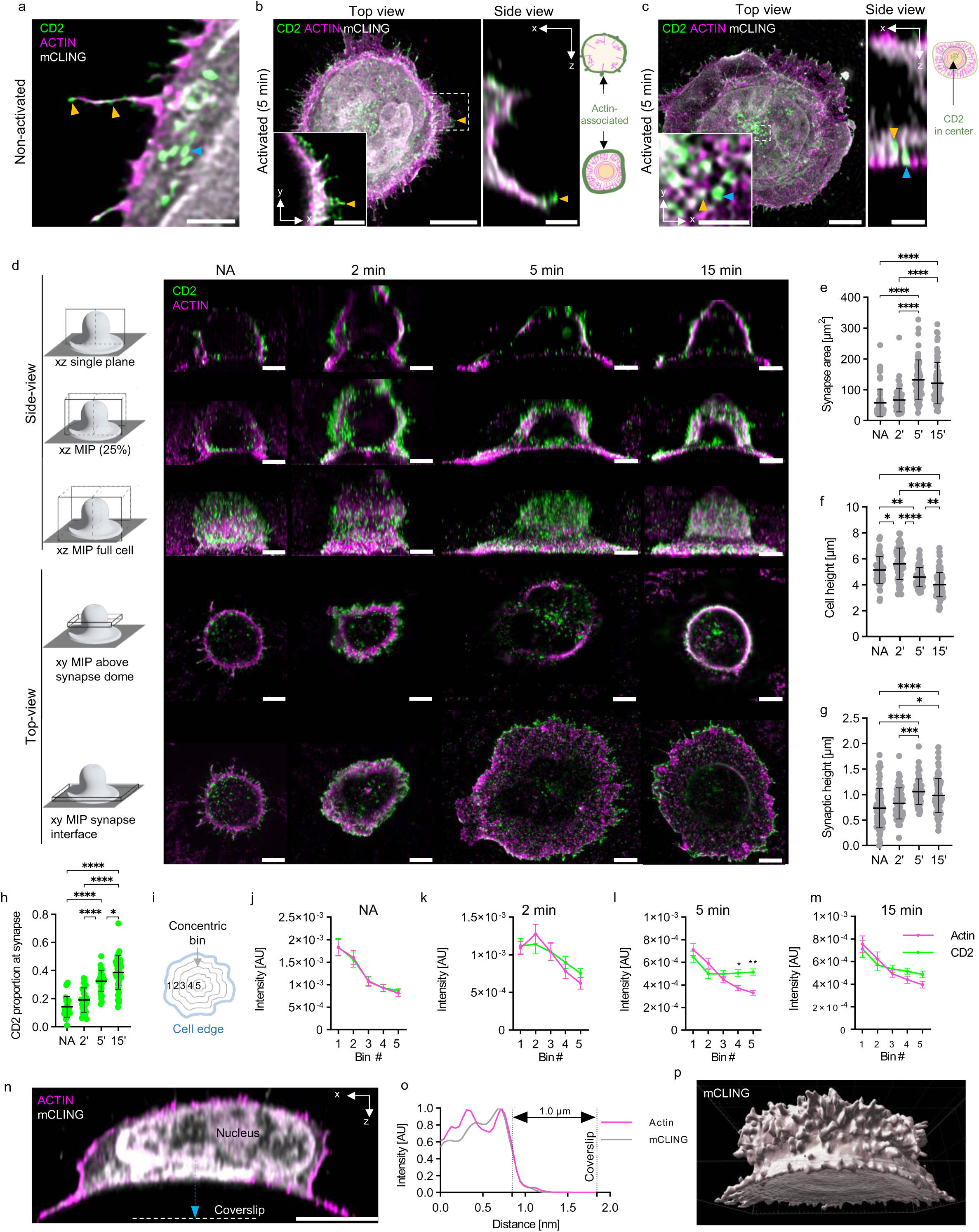
CD2 3D distribution at nanoscale. **a,** Single z-stack of confocal image of cryo-fixed and expanded CD4 T cell seeded on non-activating surface and stained for actin (magenta), CD2 (green) and mCLING (grey) to illustrate CD2 and actin associated to microvilli (orange arrowheads) and cytoplasmic CD2 localization (blue arrowhead). Scale bar: 2 µm **b ,c,** confocal image of CD4 T cell seeded on activating surface for 5 minutes and stained for actin (magenta), CD2 (green) and mCLING (grey) to illustrate CD2 distributions. Top view (xy axis) of the synapse interface is shown as MIP and inset of the indicated area as single z-stack. Side view (xz axis) of the inset is shown as single stack. **(b)** shows actin-associated CD2 (orange arrowhead) at the immune synapse edge. **(c)** illustrates CD2-rich nano-clusters at the immune synapse center, localizing at cell membrane (blue arrowhead) or within the cytoplasm (orange arrowhead). Schematic representation of CD2 and actin location is presented below, scale bar: 500 nm. **d,** cryo-ExM of CD4 T cells seeded on non-activating (NA) or activating surface for 2, 5 and 15 minutes and labeled for actin (magenta) and CD2 (green). (left) Scheme illustrating the 3D image processing of the widefield fluorescent images displayed. The first 3 rows represent side-views (xz axis) of either single planes (top row), max intensity projections (MIP) of 25% of the volume (2^nd^ row) or the full volume (3^rd^ row). The lower two rows show MIP top views (xy axis) above the dome region (∼ 5% of the full z stack volume), or of the synapse interface (between the contact region with coverslip to the synaptic dome). Within each individual time point, the same cell is displayed, scale: 2µm. **e-g,** measurement of the synaptic area (**e**), cell height (**f**) and synaptic height (**g**) for CD4 T cell at each activation time points presented in (**d**). Each dot represents one cell. Data from two healthy donors. Kruskal-Wallis with Dunn’s correction, mean ± SD is shown in (**e**). One-way Anova with tuckey’s correction, mean ± SD is shown in (**f, g**). **h**, Quantification of CD2 fluorescence intensity ratio between the synapse interface (from coverslip to synaptic dome) and the entire CD4 T cell (from coverslip to the top of the cell) at indicated time points. Each dot resembles one cell. Data from two healthy donors. One-way Anova with tuckey’s correction, mean ± SD is shown. **i,** scheme of five concentric rings used as bins to measure actin and CD2 mean fluorescence intensities at indicated activation time point presented in (e). **j-m,** quantification of mean fluorescence intensity (arbitrary unite) per bin in non-activated (**j**) cell and after 2 min (**k**), 5 min (**l**) and 15 min (**m**) of activation. Data from two healthy donors. Two-way Anova with Šídák’s multiple comparisons test, mean ± SD is shown. **n**, single plan confocal cryo-ExM image of CD4 T cell (xz, side view) seeded on activating surface for 5 min and labeled for actin (magenta) and mCLING (grey). The white dashed line marks the coverslip surface. Scale bar: 2 µm. **o**, Plot profile of normalized mCLING (grey) and actin (magenta) pixel intensity (arbitrary unite) along the dashed blue arrow (nm) in (o). The measured distance between the coverslip and the synaptic dome (50% of the fluorescence intensity) is shown. **p**, 3D surface reconstruction of the mCLING signal in (o) showing the synaptic dome. Statistical tests are indicated, ns: non-significant, * p<0.05, ** p<0.01, **** p<0.0001.

In addition, we identify a previously unreported “dome-like” space between the surface and the synaptic membrane (**Fig. 3d, 3n-p**), which increases in height over time up to 1 ± 0.25 µm (**Fig. 3d, g**). This unique T cell feature may reflect the dynamic interplay between reduced adhesion, due to the absence of CD58 and ICAM-1 on the activating surface, and actin-driven centripetal forces upon TCR engagement, highlighting a potential new aspect of T cell activation. Although this has never been described in T cells on activated surfaces, it resembles a space detected by electron microscopy in what was coined stage 4 of synapses between CD4 T cells and antigen presenting cells^51^.

We next mapped Lck localization, together with actin in resting and activated primary CD4 T cells (**Extended Data** Fig. 3c-j). Total Lck is distributed on microvilli in resting T cells (**Extended Data** Fig. 3c) and towards the base of microvilli in activated T cells (**Extended Data** Fig. 3d), as previously shown^48^. We furthermore detected an accumulation of Lck at the IS center over time, corresponding to internalization of Lck containing lipid rafts and signaling termination^52^ (**Extended Data** Fig. 3f-j). Overall, these findings highlight 3D details of T cell activation and immune synapse formation, providing new insights into the spatiotemporal organization of key adhesion and signaling molecules.

CD8 T cells release lytic molecules at immunological synapses to eliminate malignant or infected cells^53^. These cells polarize lytic granules, surrounded by LAMP1^54^ and containing Perforin (Perf), Granzyme B (GrzB)^55^, and supra-molecular attack particles (SMAPs) towards the immune synapse^56, 57^. The composition of these granules can vary among cytolytic CD8 T cells based on their antiviral activity^58^, yet their structure, quantity, shape, and distribution across different T cell subsets are largely unexplored due to their small size. Early electron microscopy studies suggested around 25 granules per cell, each about 700 nm in diameter in human and mouse CTLs^59, 60^. However, a 2013 study using patch clamp and TIRF microscopy found that lytic granules are about 418 nm in diameter at the moment of fusion with the synaptic plasma membrane^56^. The imaging improvement provided by Cryo-ExM enables not only the preservation and visualization of lytic granules (**Fig. 4a)** but also the monitoring of their structure and content loading (**Fig. 4b-e, Extended Data Video 2)**, which may vary depending on immune cell type or activation state^10^. Co-staining of Perf and GrzB enable visualizing their distribution within the LAMP1 vesicles, whether being present simultaneously or separately.

**Figure 4:**
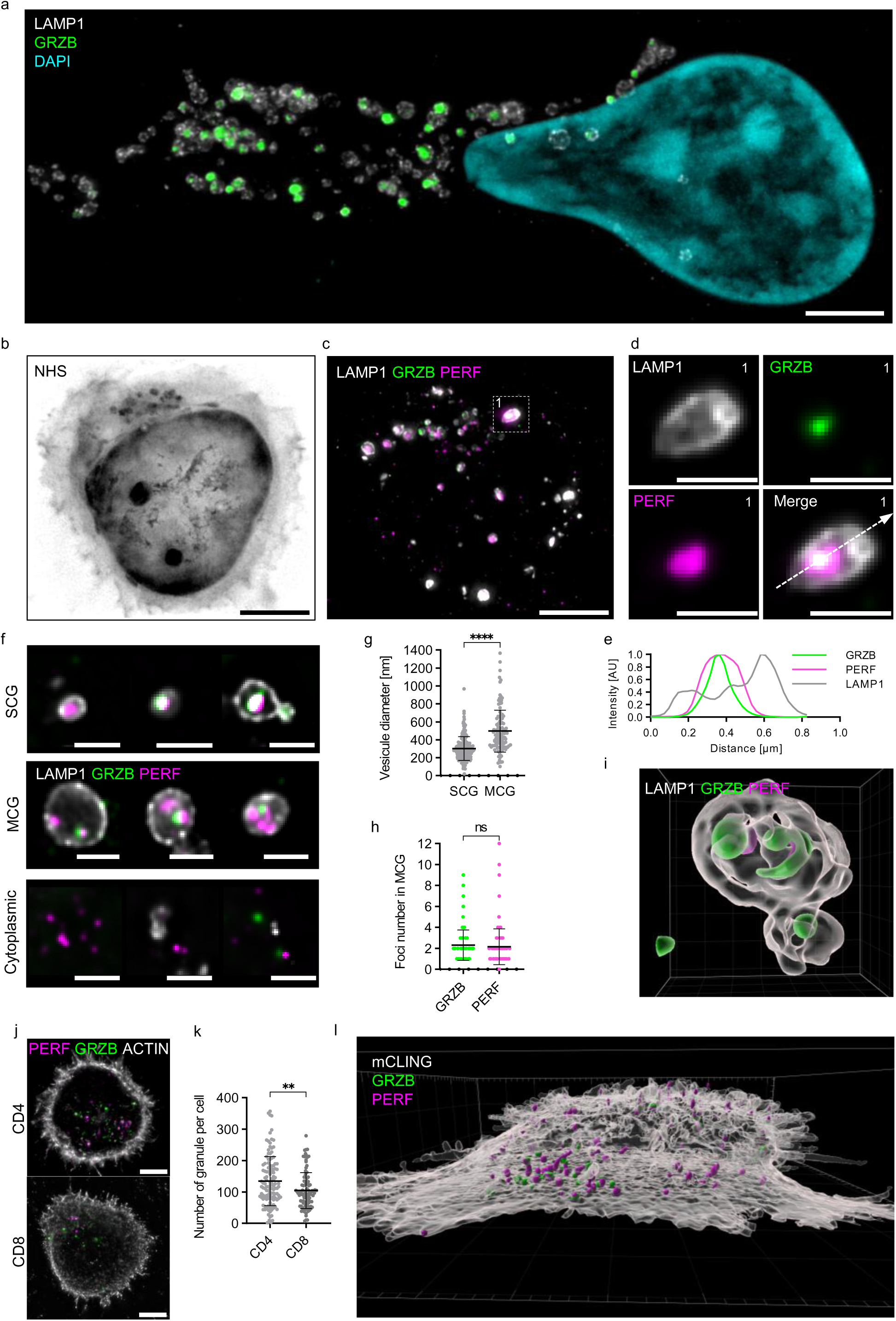
Cryo-ExM reveals the nanoscale molecular organization of T cells’ lytic granules. **a,** Representative widefield image (MIP) of cryo-fixed, expanded primary T cell labelled for LAMP1 (grey), Granzyme B (GRZB, green) and nucleus (DAPI, cyan). Lytic granules are distributed throughout the cytoplasm and the uropod of a migrating T cell, scale bar: 2 µm. **b-d,** representative widefield fluorescent images of a CD8 T cells plated on activating surface for 5 min before undergoing Cryo-ExM. The cells were stained for NHS (inverted greyscale, shown in b), LAMP1 (grey), Granzyme B (GRZB, green), Perforin (Perf, magenta), as presented in (c). MIP images are shown. **d**, inset 1 from (c) showing one lytic granule in single colors for LAMP1, GrzB and PERF or merge channels. The white dashed arrow indicates the position of the plot profile (e). b, c, scale bar: 2 µm, d, scale bar: 500 nm. **e,** Plot profile of normalized LAMP1 (grey), PERF (magenta) amd GRZB (green) pixel intensity (arbitrary unite) along the dashed white arrow (nm) in (d). **f,** representative widefield images (MIP) of CD8 T cell single core (SCG), multicore (MCG), and cytoplasmic lytic granules containing one or multiple GRZB (green) and PERF (magenta) foci surrounded by LAMP1 (grey) or not (cytoplasmic), scale bar: 500 nm. **g,** quantification of CD8 T cell SCG and MCG vesicle diameter from (f). Data from two independent donors. Each dot represents one LAMP1 positive vesicle. Mann-Whitney test, mean ± SD is shown. **h**, quantification of CD8 T cell GRZB and PERF foci number within individual MCG from (f). Each dot represents one MCG. Mann-Whitney test, mean ± SD is shown. **i**, 3D surface reconstruction of one SCG and one MCG from CD8 T cell. Lamp1 (transparent white), Granzyme B (GrzB, green), Perforin (PERF, magenta). **j**, representative widefield image (MIP) from Cryo-ExM of non-activated CD4 and CD8 T cell stained for actin (grey), Granzyme B (green) and Perforin (magenta), scale bar: 2 µm. **k**, quantification of the total number of lytic foci (Granzyme B + Perforin foci) per cell. Each dot represents one single cell. Mann-Whitney test, mean ± SD is shown. **l**, 3D surfaces reconstruction of a confocal image showing a CD8 T cell seeded on activating surface for 5min before undergoing Cryo-ExM, and labeled for mCLING (transparent grey), Granzyme B (green) and Perforin (magenta). White arrow heads point onto T cell microvilli. Statistical tests are indicated, ns: non-significant, ** p< 0.01, **** p<0.0001.

In 2020, supramolecular attack particles (SMAPs) were identified, distinguished by their glycoprotein shell instead of a traditional lipid membrane, and can independently kill targets, presenting new cancer therapies^57^. Crucial to understanding their origin, a 2022 study using mass spectrometry and electron microscopy showed that in mice, multi-core lytic granules (MCGs) develop into SMAPs, yet this has not been observed in humans^61^. Here, cryo-ExM reveals the presence of MCGs in primary human CD8 T cells (**Fig. 4f, Extended Data Video 3**) and allows detailed measurements of their size and content. Lytic proteins are organized either in single core granules (SCG), MCG or can be seen in the cytoplasm without surrounding LAMP1 (**Fig. 4f**). The dimensions of SCGs and MCGs in cryo-expanded cells align with earlier electron microscopy findings^61^ (**Fig. 4g**). Most MCGs contain 2 units of GrzB and Perf, though some carry up to 12 within one LAMP1 shell (**Fig. 4h**). Cryo-ExM thus enhances our understanding of SMAP and lytic granule biogenesis, a role previously filled by electron microscopy alone.

To quantify the structure, content, and abundance of lytic entities within 3D space (**Fig. 4i**, **Extended Data Videos 3, 4**), we developed a machine learning-supported bioimage analysis pipeline for automatic detection and 3D segmentation of lytic entities (**Extended Data** Fig. 4a). This pipeline assesses content, volume, derived diameter, position, and abundance of GrzB- and Perf-positive entities, considering the 3D volume of individual cells (**Extended Data** Fig. 4b-c). By comparing nanoscale lytic entity profiles between primary CD4 and CD8 T cells from the same healthy donor and at the same cell age, we found that while CD8 T cells are recognized for cytolytic activity, CD4 T cells, although less typically cytolytic, contain more GrzB and Perf positive lytic entities per cell (**Fig. 4j-k**). Analysis shows a higher prevalence of Perf-positive entities in CD4 T cells (**Extended Data** Fig. 4d-e), and both T cell types have slightly more Perf than GrzB positive entities (**Extended Data** Fig. 4e). CD4 cells also exhibit larger entity volumes and diameters, along with increased fluorescence intensities for Perf and GrzB (**Extended Data** Fig. 4f-h). Granule diameter distribution across CD4 and CD8 cells are comparable (**Extended Data** Fig. 4i**, 4j**), aligning with published data^60^. This data invites further investigation into the physiological implications of these granule content differences. We next investigated the granule 3D position within the cell, with respect to the IS and to the dome (**Fig. 4l**, **Extended Data** Fig. 4k-o). Both CD4 and CD8 show a significant proportion of granules that localized near the synaptic surface 5 minutes after T cell seeding on activated surfaces (**Fig. 4l, Extended Data** Fig. 4 **n-o, Extended Data Video 8**).

Since the cytotoxic function of T cells rely on its killing ability once recognizing a tumor cell, we next focused on the visualization of the molecular organization at the synaptic cleft between a T cell and its target (**Fig. 5**). We generated TCR-transgenic T cells using a previously described HLA-A2 restricted TCR recognizing the common tumor associated antigen survivin^62^. Upon co-culture of TCR-T cells with HLA-A2+survivin+ BV173 cells, a B cell precursor leukemia cell line,^62^ (**Fig. 5a**), cells were cryo-fixed and stained for NHS-ester, to visualize the cellular content (**Fig. 5a, c**) as well as actin and tubulin (**Fig.5b, 5d**). NHS-ester staining unveils the synaptic cleft formed at the T cell and tumor target intersection; a feature mostly attainable by electron microscopy that can now be imaged with a widefield microscope (**Fig. 5c, inset**). In addition, actin depletion could be readily imaged as well as centrosome polarization towards the IS formed between the cell pair (**Fig. 5d-e, Extended Data Video 5**).

**Figure 5:**
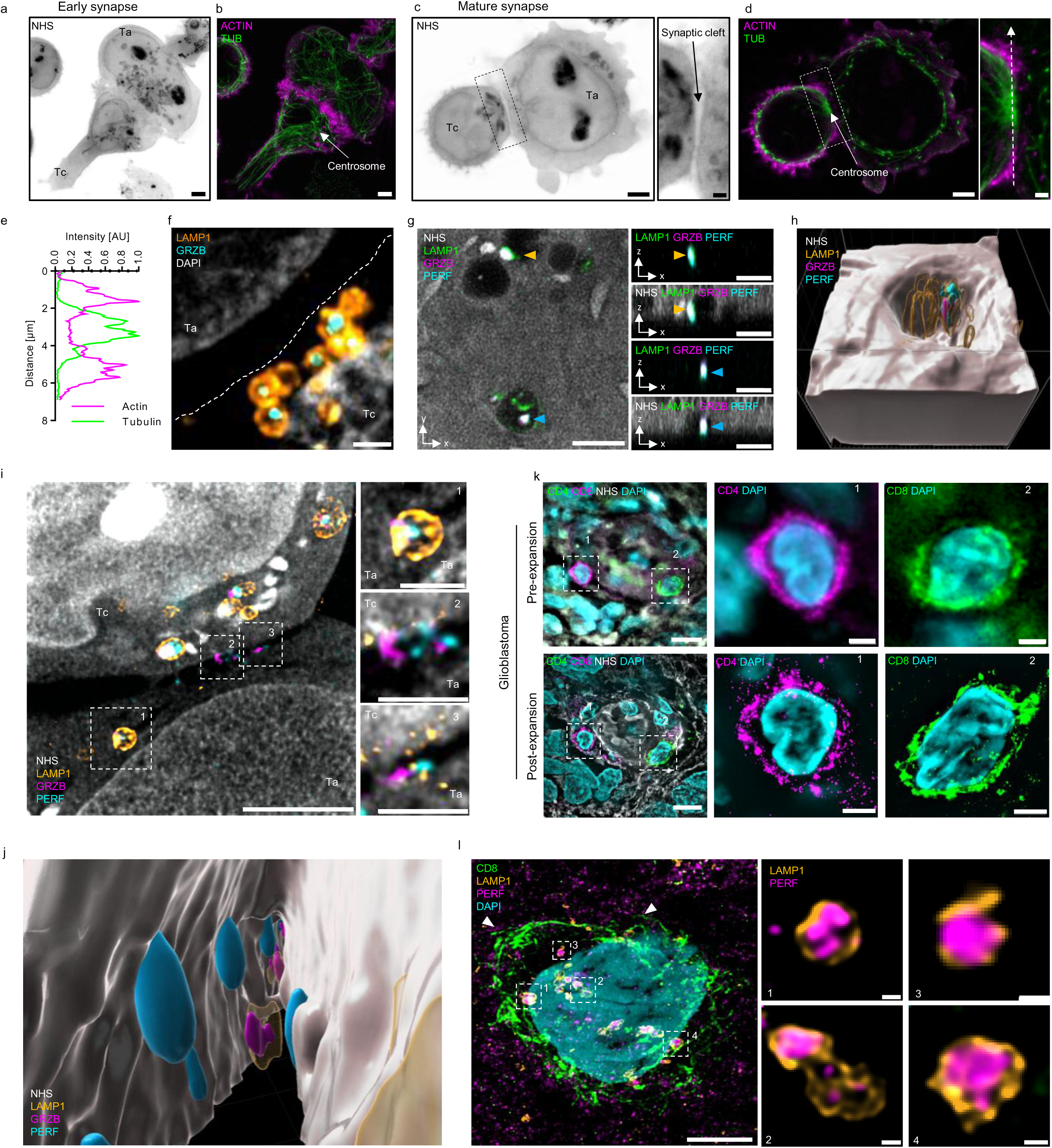
Nanostructure of TCR-engineered T cell – target cell pairs and T cells in glioblastoma tumor tissue. **a-d,** representative widefield fluorescent images (MIP) of cryo-fixed, expanded pairs of engineered TCR-transgenic (HLA-A2 survivin) human T cells (Tc) and BV173 target (Ta) cells labelled for NHS (inverted greyscale, a, c), actin (magenta, b, d) and tubulin (green, b, d). Displayed are early synapse (a, b) with the centrosome far from contact site, and mature synapse (c, d) with polarized centrosome (d). (c) inset magnifies the synaptic cleft. (d) inset magnifies centrosome polarization and actin exclusion from the synapse interface, scale 2µm. **e**, plot profile of normalized actin (magenta) and tubulin (green) pixel intensity (arbitrary unite) along the dashed white arrow (µm) in the inset image in (d). **f,** representative widefield fluorescent images (MIP) of cryo-fixed and expanded engineered TCR-transgenic human T cell and BV173 target cell pair labelled for LAMP1 (orange), Granzyme B (GRZB, blue) and nucleus (DAPI, grey). The cell-cell interface is represented by the dashed line, showing lytic granules polarized toward the target cell, scale bar: 500 nm. **g,** Single z-plane confocal images of a cryo-fixed and expanded CD8 T cell seeded on activating surface for 5min and labelled for NHS (grey), LAMP1 (green), GRZB (magenta) and Perforin (Perf, cyan). Top view (xy axis) image is shown on the left and side view (xz axis) images of the corresponding orange and blue arrowhead pointing at released lytic granules are presented on the left, scale bar: 500 nm. **h,** 3D surfaces reconstruction of the released lytic granule presented in (**g**) (blue arrowhead). NHS (grey), LAMP1 (green), GRZB (magenta) and Perforin (Perf, cyan). **i,** confocal images (MIP) of a cryo-fixed and expanded TCR-engineered (anti-survivin) human T cell (Tc) and BV173 target cell (Ta) pair labelled for NHS (grey), LAMP1 (orange), Granzyme B (magenta) and Perforin (cyan). Insets one, two and three magnify LAMP1 vesicle containing Granzyme B and Perforin in the target cell (inset 1), and free Granzyme B and Perforin within the synaptic cleft (inset 2 and 3), left scale bar: 2 µm, inset scale bar: 500 nm. **j**, 3D surface reconstruction of the confocal image presented in (**i**), showing Granzyme B (magenta), Perforin (blue) and LAMP1 (orange) within the synaptic cleft (NHS, grey). **k-l**, Immunofluorescent labeling of human glioblastoma tissue from FFPE sections. **k**, widefield fluorescent image (MIP) of a pre-expansion (upper row) FFPE section stained for CD4 (green), CD8 (magenta), nucleus (DAPI, cyan) and NHS (gray) and post-expansion (lower row) of the same sample that was re-labeled for the same cell markers. The same field of view is presented for the merge channel image and the insets 1 and 2. Individual CD4 and CD8 T cells can be detected, scale bar: 2 µm. **l**, confocal image (MIP) of expanded FFPE glioblastoma section labeled for CD8 (green), LAMP1 (orange), Perforin (magenta) and nucleus (Dapi, cyan). Insets 1-4 show single core and multicore granules within a CD8 T cell, left scale bar: 2 µm, inset scale bar: 100 nm.

Next, we focused on the lytic granules and their content at the cell-cell contact side (**Fig. 5f**). Lytic granules deliver their content into the synaptic cleft through exocytosis, which involves the fusion of the granule membrane with the plasma membrane and subsequent release of granule content^63^. This process, crucial to immune response, remains poorly understood due to visualization challenges. We investigated release events in surface-activated T cells (**Fig. 5g-h**), as well as T cell – tumor cell pairs (**Fig. 5i-j**), observing the release of Perf and GrzB from vesicles. We detect Perf and GrzB lytic entities surrounded by Lamp1 (**Fig. 5f, i**) as well as without Lamp1 (**Fig. 5g, i**), displaying Perf and GrzB release from granules into the synaptic cleft (**Fig. 5i-j, Extended Data** Fig. 5a-d **and Extended Data Videos 6, 7 and 8**) to our best knowledge, a first in the field.

Finally, we sought to bridge the gap between *in vitro* cell analysis and *in situ* examination of pathological samples by exploring U-ExM’s application potential. Archived formalin-fixed, paraffin-embedded (FFPE) tissues in clinical pathology departments hold crucial molecular insights into human physiology and disease. While single-cell T cell analysis provides essential data on T cell biology, understanding T cell interactions within tumor tissues is vital for cancer immunotherapy. However, epifluorescence imaging of these tissues does not reveal detailed cellular structures^64^, and super-resolution microscopy encounters significant technical hurdles in tissue^65^. We hypothesize that expansion microscopy (U-ExM) of FFPE tissues could provide nanoscale structural insights into T cell biology. To test this, we adopted a tissue-specific U-ExM protocol^66, 67^ that enables the identification of specific regions within tissue sections and supports multiple rounds of post-expansion immunofluorescence staining and stripping (**Extended Data** Fig. 5e). As proof of concept, we imaged primary brain tumor tissues (**Fig. 5k-l, Extended Data** Fig. 5f), successfully identifying T cells infiltrating human glioblastoma. This approach allowed us to extract nanoscale details, such as the structural composition of T cell lytic granules (**Fig. 5l**) and CD8 T cell microvilli (**Fig. 5l, Extended Data** Fig. 5f, white arrowheads**, Extended Data Video 9**). This novel protocol provides a powerful solution for multiplexed nanoscale structural analysis of FFPE tissues.

## Discussion

Ultrastructure Expansion Microscopy, especially in combination with cryo-fixation (Cryo-ExM) emerges in this study as a powerful tool for visualizing the high-resolution near-native nanoscale architecture of T cells and their interactions with cancer cells at the immune synapse. Combining cryo-fixation with isotropic sample expansion and pan-labelling, Cryo-ExM preserves fragile structures like cytoskeletons, tunneling nanotubes, centrioles, and lytic granules. It enables detailed 3D imaging of immune synapse formation, cytoskeletal dynamics, and protein localization, including the release of cytotoxic molecules. The method is further extended to FFPE tumor tissues, offering nanoscale resolution in clinical samples, thus bridging single-cell immunology with human pathology.

## Material and Methods

### Cell culture and T cell activation

#### Cell lines

Homo sapiens bone osteosarcoma U2OS and T lymphoblast Jurkat Clone E6-1 were purchased from the American Type Culture Collection. B cell precursor leukemia BV173 and B lymphoblast MM.1S CRL-2974 cells were a kind gift from the Arber lab. All lines were maintained according to the suppliers’ instructions. U2OS were cultured in Dulbecco’s modified Eagle’s medium and GlutaMAX, supplemented with 10% FBS and penicillin - streptomycin (100 μg/ml) at 37 °C in a humidified 5% CO2 incubator. Jurkat, BV173 and MM.1S were cultured in RPMI containing 10% FBS, 2 mM L-Glutamine, 1% Penicilin-Streptomycin) at 37 °C in a humidified 5% CO2 incubator.

### Generation of retroviral vectors and supernatant

The retroviral vector expressing the survivin-specific (s24) TCR has been previously described^62^. Transient retroviral supernatant was prepared by transfection of 293T as described^68^.

### Generation of transgenic T cells

Peripheral blood mononuclear cells (PBMCs) were isolated from buffy coats by Ficoll density gradient centrifugation (Lymphoprep, StemCell #07851). Buffy coats from deidentified healthy human volunteer blood donors were obtained from the Center of Interregional Blood Transfusion SRK Bern (Bern, Switzerland).

PBMCs were activated on plates coated with anti-CD3 (1 µg/mL, Biolegend, #317347, OKT3) and anti-CD28 (1 mg/mL, Biolegend, #302934, CD28.2) in T cell medium (RPMI + 10% FBS, 2 mM L-Glutamine, 1% Penicillin-Streptomycin) with IL-15 and IL-7 (10 ng/mL each, Miltenyi Biotec, #130-095-362 and #130-095-765).

A day before transduction, non-tissue culture-treated 24-well plates (Grener Bio One, #662102) were coated with retronectin (7 µg/mL, Takara Bio, #T100B) in PBS and incubated overnight at 4°C. Three days post-activation, retronectin was removed, plates were blocked with T cell medium (15 min, 37°C), and retroviral supernatant was centrifuged onto the coated plate (2000 g, 1 h, 32°C). The supernatant was removed, and activated T cells (0.15×10⁶ cells/mL) were added, followed by centrifugation (1000 g, 10 min, 21°C). Cells were incubated (37°C, 5% CO₂) for 3 days.

After 48–72 h, T cells were harvested and expanded in T cell medium with IL-7 and IL-15.

### Immune synapse formation on activated surfaces and Cryo-fixation

12 mm coverslips were coated with poly-D-lysin (0.1 mg/ml, Gibco, #A3890401) overnight at 4°C. The coverslips were subsequently coated with Anti-CD3 (OKT3, 1 µg/ml) and anti-CD28 (1 µg/ml) for 3h at 37°C and washed with PBS before being used. T-cells were seeded at 1.5×10^5^ cells per coverslips and activated for 2 min, 5 min and 15 min. Non activated T cells were seeded for 10 min on poly-D-lysin coated coverslips only.

The cryo-fixation procedure was performed as previously described^15^. Briefly, a thin tweezer (Dumont 5, Sigma F6521-1EA) was used to hold the 12 mm coverslips containing the sample. The remaining medium was blotted with filter paper and the coverslip was quickly plunged using a plunge freezer into liquid ethane cooled at −170 °C using liquid nitrogen^15^. Note that a homemade tweezer holder was designed to maintain the tweezer aligned to the center of the cryo-chamber containing the liquid ethane, reducing the risk of sample loss.

Coverslips were rapidly transferred to 5 ml Eppendorf tubes containing 1 ml of liquid nitrogen-chilled acetone mixed with 0.1% paraformaldehyde (PFA) and 0.02% Glutaraldehyde (GA). The samples were incubated overnight on dry ice with a 45° angle and under agitation to increase the temperature to −80 °C. The samples were subsequently incubated without dry ice for 45 min to equilibrate the temperature to *∼*0 °C. Sequential ethanol: water baths containing 0.1% PFA and 0.02% GA were used to rehydrate the samples as followed: ethanol 100%, ethanol 100%, ethanol 95%, ethanol 95%, ethanol 70%, ethanol 50%, ethanol 25%, ddH2O and PBS. Cells were stored in PBS until expansion or directly processed for immunostaining.

### Ultrastructure expansion microscopy (U-ExM)

Expansion of the cells was performed as previously described^17, 20^. Briefly, cryo-fixed cells were incubated for cross linking prevention 3h in 2% acrylamide and 1.4% formaldehyde diluted in PBS at 37 °C before gelation in monomer solution (19% sodium acrylate, 0.1% bis-acrylamide, and 10% acrylamide) supplemented with 0.5% of TMED and APS. The samples were incubated on ice for 5 min followed by 1 h at 37 °C. After gelation, denaturation was performed for 1.5 h at 95 °C in denaturation buffer. Gels were washed three times for 10 min in ddH2O, allowing for expansion before the gel size was measured with a caliper to calculate the expansion factor by dividing the expanded gel size with the size of the original coverslip (12 mm).

For centriole evaluation (**Fig. 2a-f**), no pre-fixation step was performed before U-ExM procedure, allowing for cytoplasmic microtubule depolymerization. Directly after removing the cell culture medium, cells were processed for U-ExM as described above.

### Tissue expansion microscopy from FFPE samples

FFPE sections on slide were first processed for deparaffinization. The slide were heated on top of a heatblock at 60°C for 15 min before being submerged in successive baths as followed: 3 baths of 3 min in Xylene-Substitute (#A5597, Sigma-Aldrich), 2 baths of 2 min in 100% Ethanol, 1 bath of 2 min in 95% Ethanol, 1 bath of 1 min in 70% Ethanol and 2 baths of 2 min in pre-warmed (37°C) water. A spacer (IS213, Sunjin Lab) was sticked on the slide to form a chamber space around the sample. 500 µL of cross-linking prevention solution (2% acrylamide and 1.4% formaldehyde diluted in PBS) was added within the chamber space and incubated 3h at 37 °C. After removing the solution, the slide was transferred onto a cold metallic block (pre-chilled at −20°C), and 180 µL of inactivated monomer solution (19% sodium acrylate, 0.1% bis-acrylamide, and 10% acrylamide) was incubated on the sample for 15 min. The inactivated monomer solution was subsequently replaced with 400 µL of activated monomer solution (containing 0.5% of TMED and APS). The sample chamber was sealed by adding a glass coverslip on top of the spacer and the slide was left for 15 min incubation on the cold metalic block before being transferred to room temperature for 2 h.

The coverslip and de the spacer were carefully detached, and the slide was incubated in a 50 mL falcon tube containing 35 mL of denaturation buffer dure 2h at 95°C. The detached gel was then washed three times 30 min in water and allowed to fully expand.

### Authorization to work with patient’s tissue samples

Human glioblastoma samples were collected at the Geneva University Hospital in a biobanking protocol approved by the Geneva ethics committee (protocol 2022-02109). All patients provided written informed consent. Use of the biobanked samples in the current project was approved by the Geneva ethics committee under protocol 2025-00256.

### Immunolabeling

Gels were incubated in PBS for 30 min and stained for 3 h at 37 °C under constant agitation with primary antibodies (Table 1) diluted in 2% PBS–BSA. Gels were washed 3 times in PBS Tween 0.1% and subsequently stained with secondary antibodies (Table 1) diluted in 2% PBS– BSA. Gels were washed three times in PBS–Tween 0.1% and expanded by successive baths of ddH2O. When indicated, the gels were further incubated in NHS-ester diluted in PBS for 1h at 37°C under constant agitation, washed three times in PBS and expanded by successive baths of ddH2O.

**Table 1:**
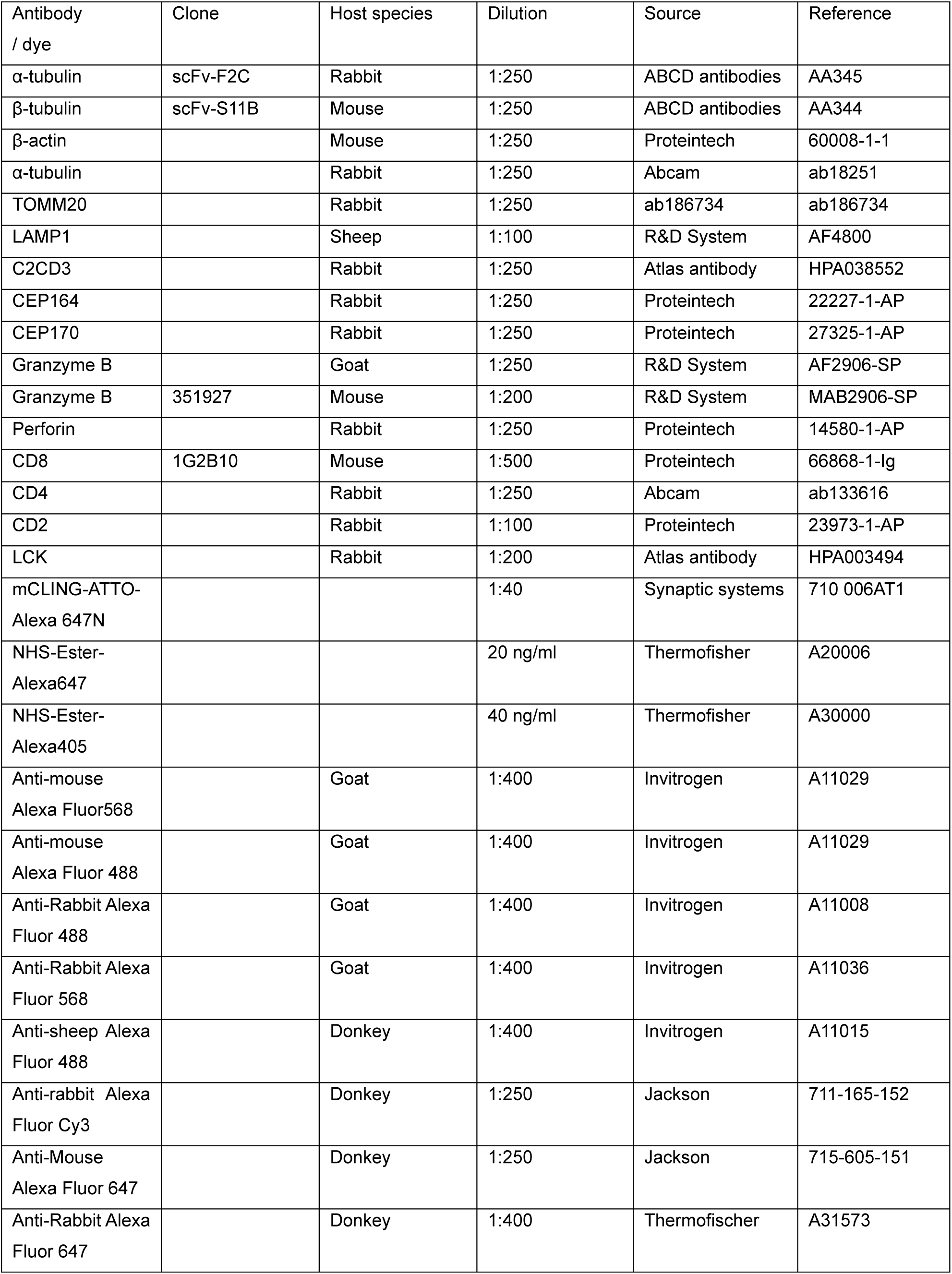
antibodies and fluorescent dyes used in this study.

| Antibody / dye | Clone | Host species | Dilution | Source | Reference |
| --- | --- | --- | --- | --- | --- |
| α-tubulin | scFv-F2C | Rabbit | 1:250 | ABCD antibodies | AA345 |
| β-tubulin | scFv-S11B | Mouse | 1:250 | ABCD antibodies | AA344 |
| β-actin | | Mouse | 1:250 | Proteintech | 60008-1-1 |
| α-tubulin | | Rabbit | 1:250 | Abcam | ab18251 |
| TOMM20 |  | Rabbit | 1:250 | ab186734 | ab186734 |
| LAMP1 |  | Sheep | 1:100 | R&D System | AF4800 |
| C2CD3 |  | Rabbit | 1:250 | Atlas antibody | HPA038552 |
| CEP164 |  | Rabbit | 1:250 | Proteintech | 22227-1-AP |
| CEP170 |  | Rabbit | 1:250 | Proteintech | 27325-1-AP |
| Granzyme B |  | Goat | 1:250 | R&D System | AF2906-SP |
| Granzyme B | 351927 | Mouse | 1:200 | R&D System | MAB2906-SP |
| Perforin |  | Rabbit | 1:250 | Proteintech | 14580-1-AP |
| CD8 | 1G2B10 | Mouse | 1:500 | Proteintech | 66868-1-Ig |
| CD4 |  | Rabbit | 1:250 | Abcam | ab133616 |
| CD2 |  | Rabbit | 1:100 | Proteintech | 23973-1-AP |
| LCK |  | Rabbit | 1:200 | Atlas antibody | HPA003494 |
| mCLING-ATTO-Alexa 647N |  |  | 1:40 | Synaptic systems | 710 006AT1 |
| NHS-Ester-Alexa647 |  |  | 20 ng/ml | Thermofisher | A20006 |
| NHS-Ester-Alexa405 |  |  | 40 ng/ml | Thermofisher | A30000 |
| Anti-mouse Alexa Fluor568 |  | Goat | 1:400 | Invitrogen | A11029 |
| Anti-mouse Alexa Fluor 488 |  | Goat | 1:400 | Invitrogen | A11029 |
| Anti-Rabbit Alexa Fluor 488 |  | Goat | 1:400 | Invitrogen | A11008 |
| Anti-Rabbit Alexa Fluor 568 |  | Goat | 1:400 | Invitrogen | A11036 |
| Anti-sheep Alexa Fluor 488 |  | Donkey | 1:400 | Invitrogen | A11015 |
| Anti-rabbit Alexa Fluor Cy3 |  | Donkey | 1:250 | Jackson | 711-165-152 |
| Anti-Mouse Alexa Fluor 647 |  | Donkey | 1:250 | Jackson | 715-605-151 |
| Anti-Rabbit Alexa Fluor 647 |  | Donkey | 1:400 | Thermofischer | A31573 |

For direct immunolabeling (**Extended Data** Fig. 1a, b) coverslips were incubated for 15 min in 2% PBS–BSA–Tween 0.1% and incubated with the primary antibodies diluted in 2% PBS– BSA–Tween 0.1% for 2 h at room temperature. After three washes in PBS–Tween 0.1%, secondary antibodies diluted in 2% PBS–BSA–Tween 0.1% were incubated for 1 h at room temperature before being washed three times with PBS-Tween 0.1%. Nuclei were stained with 4,6-diamidino-2-phenylindole (DAPI) in PBS for 5 min at room temperature. Coverslips were then mounted on glace slides and imaged as described below. Primary and secondary antibodies were diluted twice compared to post-expansion staining.

### Image acquisition

Pieces of gels were mounted on 24 mm round precision coverslips (1.5H, #0117640, Marienfeld) coated with poly-D-lysine for imaging. Image acquisition was performed using a 63x 1.4 NA oil immersion objective with an inverted widefield Leica DMi8 Thunder microscope or a 63x 1.4 NA water immersion objective with a confocal Leica Stellaris 8 FALCON microscope. The microscope parameters were controlled using the LAS X software (LAS X; Leica Microsystems). In the figures, widefield images were treated with large volume computational clearing (LVCC) mode at max resolution, adaptative “strategy” and water as “mounting medium” (Thunder, Leica Microsystem), and confocal stack images with lightning noise reduction (Leica Microsystem).

Widefield three-dimensional stacks images (Leica DMi8 Thunder) were acquired with a voxel size of 0.1013 x 0.1013 x 0.2131 µm and confocal three-dimensional stack images (Leica Stellaris 8 FALCON) were acquired in Lightening mode with adapted voxel size. Image denoising is performed immediately through the Leica Thunder software.

### Quantifications

Each measurement was divided by the calculated expansion factor of the corresponding gel and reported as such in the graphs or figure scale bars. Unless specified, denoised images were used for representation and quantifications.

### Centriole measurements

Analysis on centrioles was performed on unfixed cells and images were analyzed using Fiji. Measurement of the centriole length and diameter was performed on side views using tubulin staining as previously described^69^.The pixel size of the image was artificially divided by 6 to improve the accuracy (plugin “CropAndResize”, https://github.com/CentrioleLab) before measuring the fluorescent signal distribution of tubulin using the Fiji line scan (with a width of 200-220 pixels) and the plot profile tool. Using the plugin “PickCentrioleDim” (https://github.com/CentrioleLab) to facilitate the picking of the start and the end of the fluorescent signal (defined as 50% of the peak value at both extremities of the centriole), the user automatically generates an entry table with the coordinates of fluorescent signal extremities of the tubulin channel.

The coordinates of fluorescent signal extremities of the tubulin were used to calculate the distance between two centriole extremities and plotted using GraphPad Prism 9.

### Extraction of electron-microscopy images from OpenOrganelle

Scanning electron microscopy data used in Extended Data Figures 2a, b was downloaded from OpenOrganelle website: https://openorganelle.janelia.org/datasets (datasets jrc_jurkat-1, jrc_ctl-id8-2, jrc_ctl-id8-1) ^34^. The 3D view was obtained with the UCSF chimera software. Top views of the centrioles were generated using ImageJ software and the “reslice” function to reorient the centrioles and obtain an adequate view.

### Imaris segmentation and representation

The cell 3D surface reconstructions and video editing were performed using bitplane Imaris 10.1 software. After background subtraction for all channels, volumetric surfaces of channels of interest were reconstructed using the trainable pixel classifier algorithm.

### Nucleus dimension for isotropy evaluation

Primary T cells and Jurkat cells were seeded at 3.5 x 10^5^ cells on 12 mm coverslips coated with poly-D-Lysine, OKT3 (1 µg/ml) and anti-CD28 (1 µg/ml) and incubated 10 min. Cells were cryo-fixed as described above and were incubated for crosslinking prevention 3h at 37°C as described in the U-ExM procedure. Cells were subsequently washed three times with PBS and stained with DAPI (1 ng/ml) in PBS for 5min. After three washes in PBS, the coverslips were placed on a QR chamber for 12 mm coverslips (Warner instruments, QR-48LP, 64-1946) with PBS and imaged with a 63x 1.4 oil immersion objective using the widefield Leica DMi8 Thunder microscope. Cells were subsequently processed according to the U-ExM protocol as previously described^69^. Cells were stained a second time with DAPI (1 ng/ml) in PBS for 15 min before being washed 3 times with ddH_2_O, allowing for expansion. Pieces of expanded gels were imaged for DAPI with the same microscope parameters used to image non expanded cells. Using automatic segmentation of the DAPI-stained nuclei, the Nuclear Cross Section (NCS) value, defined as the square root of the nucleus areas, was determined for both non-expanded and expanded cells (divided by the gel expansion factor) as previously described^70^ using Fiji.

### Synaptic height, cellular height and synaptic area

The synaptic height and the cell size were measured on expanded non-activated and activated primary T cell stained for actin. The synaptic height was calculated as the distance between the z coordinates of the stack where the first actin signal appears at the glass interface (referred to as synaptic plane) and the z coordinate where the actin signal of the dome (referred to as synaptic dome) disappears multiplied by the voxel depth (0.2131 µm).

The cell height was measured as the distance between the z-coordinates of the synaptic plan and the top of the cell multiplied by the voxel depth.

To measure the synaptic area, the cell edges were first automatically defined based on actin and DAPI channels using the U-ExM-synapse-segmentation macro (https://github.com/CentrioleLab) and corrected manually when necessary. The synaptic area of each cell was determined by measuring the surface within the cell edges.

### Concentric bin analysis and protein repartition at synapse

Images were analyzed using homemade Fiji macros. The cell edges were first automatically defined based on Actin and DAPI channels using the U-ExM-synapse-segmentation macro (https://github.com/CentrioleLab) and corrected manually when necessary. The z coordinates of the synaptic region were defined manually using the orthogonal viewer to perform a sum z-stack projection. The cell contour was used as a region of interest of reference and scaled down 5 times to create centered concentric bins. The fluorescence raw integrated density was measured within each concentric bin for Actin, CD2 and LCK staining.

To evaluate the total proportion of CD2 and LCK protein at the synapse region, the homemade synapse protein intensity Fiji macro was used (https://github.com/CentrioleLab). Fluorescence raw integrated density of the sum z-stack projection of the synaptic region was measured and reported to the fluorescence raw integrated density measured on the sum z-stack projection of the entire cell. All measurements were corrected for background intensity and assembled using R environment before being plotted with GraphPad Prism 9.

### 3D granule segmentation

For 3D segmentation of Perforin and Granzyme B (GrzB) lytic granules, we used the machine learning-based LABKIT ^71^ plugin in Fiji to create pixel classifiers for each marker. Training datasets were prepared from eight randomly selected 3D image stacks (31 slices each) from three independent experiments. These datasets, separated by channel, were loaded into LABKIT, and the following 3D features were selected: Sigma (1.0, 2.0, 5.0, 8.0), Gaussian blur, difference of Gaussians, gradient magnitude, Laplacian of Gaussian, Hessian eigenvalues, structure tensor eigenvalues, mean filter, and variance filter. Granules were manually annotated as foreground (Perf or GrzB signal) or background, and the classifiers were trained accordingly. When necessary, labelled images were manually refined, and classifiers were retrained for improved accuracy. The trained classifiers were then applied within a custom Fiji macro for segmentation of lytic granules in primary T cells.

Cell edge detection was performed automatically using a Fiji macro. The orthogonal slicer function was used to determine the z-coordinates of the synaptic plane, synaptic dome, and the top of the cell. Total fluorescence intensity for Perforin and GrzB was measured using maximum intensity z-projections from the synaptic plane to the top of the cell. The trained classifiers segmented granules in 3D, generating probability maps that were converted into 8-bit images and thresholded, followed by application of the Otsu algorithm to generate binary masks. The 3D Object Counter in Fiji was used to filter objects smaller than 12 voxels (84 nm diameter), generating object maps for each cell. The 3D Object Manager from the 3D Suite plugin was used for metric and fluorescence intensity measurements of individual granules, with corrections for background fluorescence.

To assess granule colocalization, 3D MultiColoc in Fiji was used to calculate overlapping volumes. The total granule count per cell was determined by summing Perforin and GrzB granules and subtracting colocalized granules. The polarization index of each granule was calculated as the distance from the synaptic dome to the granule center of mass, multiplied by voxel depth. Data analysis was performed in R, and plots were generated using GraphPad Prism 9.

### Vesicular measurement

Several LAMP1 positive vesicles per cells were cropped in x, y and z dimension and resized by artificially dividing the pixel size by 6 to increase measurement accuracy using a homemade Fiji macro (https://github.com/CentrioleLab). LAMP1 vesicle diameter was measured using the homemade Fiji macro (https://github.com/CentrioleLab) that was adapted from the “PickCentrioleDim” plugin. The Fiji line scan and the plot profile tool were used to determine the wider diameter of the vesicle (50% of the peak value at both extremities). Then, the number of Perforin and GranzymeB positive granules within LAMP1 vesicles were counted manually using the Multipoint tool in Fiji.

### Statistical analysis

Images were analyzed using Fiji. Statistical analyses were performed using GraphPad Prism 9, as indicated below with details and are provided in the figure legends and results. All data are expressed as the mean (average) +/− the standard deviation (SD) or the median for logarithmic scales. The sample sizes (n) and what they represent (e.g., number of granules, number of replicates) are stated in figure legends and/or results. Normal distribution was systematically tested. Parametrical tests were performed when the data were normally distributed, while non-parametrical test were used if not. Applied statistical tests are indicated in the figure legends.

### Data availability

The data that support the findings of this study are available as ‘source data’ provided with the manuscript. Further request can be sent to the corresponding authors.

## Author contributions

BW and VH designed the study. BW and FL planned, performed and analyzed experiments. BW, VH, PG and FL interpreted the data. BW, VH and FL wrote the manuscript. PG contributed to and discussed the manuscript. OM and IM established the tissue expansion, performed tissue expansion experiments and collected and analyzed tissue expansion data. FL wrote code in Fiji and R for image analysis and created IMARIS 3D renderings. E.P. carried experiments and designed analysis pipelines. VD and DM contributed patient material and corrected the manuscript. JR contributed to vector cloning and T cell transductions. CA advised on T cell transduction, corrected the manuscript.

## Acknowledgement

We thank Daniel Sage (EPFL) for advice on the bioimage analysis. We thank Sónia Gomes Pereira for extraction of electron microscopy images from Openorganelle.janelia.org. We thank Claire Hivroz (Institute Curie) for her comments and input on the manuscript. This work was supported by the ISREC TANDEM foundation attributed to V.H. and B.W., the Stiftung für Krebsbekämpfung CH to B.W., the Swiss National Science Foundation (SNSF) 310030_205087 attributed to P.G and V.H and the Gelbert foundation attributed to VH.

## Declaration of interest

C.A. holds patents and provisional patent applications in the field of engineered T cell therapies. C.A. receives licensing fees and royalties from Immatics (through previous institution Baylor College of Medicine), participated in advisory boards for Kite/ Gilead, Janssen and Celgene/ BMS, received sponsored travel from Gilead (through current institution University Hospital Lausanne). D.M. is an inventor of patents related to CAR-T cell therapy, filed by the University of Pennsylvania, the Istituto Oncologico della Svizzera Italiana (IOSI), and the University of Geneva. D.M. is a consultant for Limula SA, and MPC Therapeutics SA., DM is the scientific co-founder and has an equity interest in Cellula Therapeutics SA. All other authors declare no competing interests.

## Extended Data Fig. Figure legends

**Extended Data Figure 1 Cellular nanostructure preserved by cryo-freezing**

**a,b**, PFA fixed (a) or cryo-fixed primary human T cells (b) were stained for actin (magenta), α-tubulin (green) and nucleus (Dapi, cyan), and imaged with a widefield microscope, scale: 5µm. **c,d**, representative widefield image (MIP) of a cryo-frozen Jurkat T cell non-expanded (c) and expanded (d), stained for actin (magenta), α-tubulin (green) and nucleus (DAPI, blue), scale bar: 5 µm.

**e**, nuclear cross section (NCS) measurement of Jurkat cells before and after undergoing cryo-ExM. Each dot represents one nucleus. Data from n = 3 experiments, corrected for expansion factor. Mann-whitney test, mean ± SD is shown, * p< 0.05.

**f-h**, cryo-fixed and expanded pairs of TCR engineered (anti-survivin) human T cells (Tc) and BV173 target (Ta) cells labelled for NHS (grey) and TOM20 (green). **f**, representative widefield fluorescent images (MIP). **g**, 3D reconstruction of the image shown in (**f**), showing the fluorescent signal of NHS and TOM20. **h**, 3D surface reconstruction of the cell surface (transparent grey). Mitochondria surfaces were segmented using NHS (magenta) and TOM20 (transparent green) florescent signal. Inset 1 is presented in (i).

**Extended Data Figure 2 T cell centriolar architecture and IS maturation stages**

**a, b,** Electron microscopy images (https://openorganelle.janelia.org/) of Jurkat (a) and primary human T cells (b), scale bar: 100 nm.

**c,** representative westernblot of full cell lysates from U2OS, primary CD4 and CD8 T cells and Jurkat cells stained for CEP170 (170kDa) and actin as a loading control (40kDa).

Representative form two experimental replicates of 2 healthy donors.

**d, e,** representative immunofluorescence images of PFA fixed, non-expanded Jurkat (d) and primary human CD4 T cells (e) plated on non-activating (NA) or activating surface for 2, 5 and 15 min. Cells were stained for actin (magenta), α-tubulin (green) and nucleus (DAPI, blue). Upper row: low magnification, scale bar: 10 µm. Lower row: insets from above at higher magnification, scale bar: 5 µm.

**f,** Scheme showing the different cellular morphology used to classify immune synapse maturation stage into non-activated (NA), early activation (early), early ring, ring, early contraction and contraction based on the actin organization and cell morphology.

**g-i,** proportion of the different activation status for non-activated and activated Jurkat **(g)**, primary CD4 **(h)** and CD8 **(i)** T cell based on the actin organization and cell morphology classification presented in (f) at indicated timepoints. Data from n = 3 experiments for Jurkat cells, n = 1 experiment for primary CD4 and CD8 T cells. Mean and SD are shown.

**Extended Data Figure 3 Lck 3D distribution at nanoscale**

**a-b ,** representative wide-field fluorescent images of cryo-fixed and expanded CD4 T cell plated on non-activating (a) or activating (b) surface for 5 min and stained for actin (magenta) and CD2 (green). **a,** single z-stack showing cortical actin associated CD2 (low magnification). CD2 accumulates into cytoplasmic foci (arrowhead, inset 1) and at the microvilli tip (inset 3, arrowhead), scale bar: 500 nm.

**b**, single z-slice taken at increasing distance (µm) from the coverslip surface (from left to right panel) showing the Actin-associated CD2 at the synapse edge (left panel), CD2 forming corolla-like accumulation (middle panel), and CD2 foci accumulation at the center of the synapse (right panel), scale bar: 500 nm.

**c, d,** representative widefield fluorescent images (single z stack) of cryo-fixed and expanded CD4 T cell plated on non-activating (c) or activating (d) surface for 5 min, and stained for NHS (inverted greyscale), Actin (magenta) and LCK (green) to illustrate LCK distributions at the tip of microvillies (c) and LCK punctiform distribution at the synapse interface (d), scale bar: 2µm.

**e,** Cryo-ExM of CD4 T cells seeded on non-activating or activating surface for 2, 5 and 15 minutes and labeled for actin (magenta) and LCK (green). The first 3 rows represent side-views (xz axis) of either single planes (top row), max intensity projections (MIP) of 25% of the volume (2^nd^ row) or the full volume (3^rd^ row). The lower two rows show MIP top views (xy axis) above the dome region (∼ 5% of the full z stack volume), or of the synapse interface (between the contact region with coverslip to the synaptic dome). Within each individual time point, the same cell is displayed, scale bar: 2 µm.

**f-i,** quantification of LCK (green) and Actin (magenta) mean fluorescence intensity (arbitrary unite) within five concentric rings used as bin in non-activated (NA, **f**) cell and after 2 min (**g**), 5 min (**h**) and 15 min (**i**) of activation. Data from n = 2 donors. Two-way Anova with Šídák’s multiple comparisons test, mean± SD is shown.

**j**, Quantification of LCK fluorescence intensity ratio between the synapse interface (from coverslip to synaptic dome) and the entire CD4 T cell (from coverslip to the top of the cell) at indicated time points of activation. Each dot resembles one cell. Data from n = 2 donors. Kruskal-Wallis test with Dunn’s correction, mean± SD is shown. Statistical tests are indicated.

* p<0.05, ** p<0.01 *** p<0.001, **** p<0.0001.

**Extended Data Figure 4 Machine learning-supported bioimage analysis pipeline for automatic detection and 3D segmentation of lytic entities**

**a,** automated image processing pipeline using machine learning based labkit plugin and Fiji/ImageJ homemade macros to perform 3D segmentation and measurements of Granzyme B and Perforin lytic granules.

**b, c**, primary CD4 T cell plated on activating surface before undergoing cryo-ExM, and labeled for Actin, Granzyme B and Perforin and image with widefield fluorescent microscope.

b, representative examples of segmented Perforin (seg-PERF, left) and Granzyme (seg-GRZB, right) lytic particles showed as z-stack projection of a side view (xz axis, upper row) or top-view (xy axis, lower images). The color code indicates the 3D position of the entity within the z-stack.

c, same images as in (b) showing the overlay of the segmented Perforin (seg-PERF, cyan) with Perforin (magenta) fluorescent signal (left) and segmented Granzyme B (seg-GRZB, blue) with the Granzyme B (GRZB, green) fluorescent images.

**d**, representative widefield fluorescent images (MIP) of cryo-fixed and expanded primary CD4 and CD8 T cells plated on non-activating or activating surface for 5min, and labelled for Actin (grey), Granzyme B (green) and Perforin (magenta). Insets show granzyme B and Perforin lytic granule. Low magnification scale 2µm. inset scale bar: 500 nm.

**e**, Number of Granzyme B (green) and Perforin (magenta) lytic entities per CD4 and CD8 T cell. Each dot represents one cell. Kruskal-Wallis test with Dunn’s correction, mean± SD are shown.

**f**, quantification of the total granzyme B (green) and Perforin (magenta) mean fluorescence intensity (arbitrary unite) per CD4 and CD8 T cell. Each dot represents one cell. Kruskal-Wallis test with Dunn’s correction, mean± SD are shown.

**g**, measurement of the average of Perforin (magenta) and Granzyme B (green) lytic entity volume (µm^3^) for primary CD4 and CD8 T cell. Each dot represents one cell. Kruskal-Wallis test with Dunn’s correction, mean± SD are shown.

**h**, quantification of Perforin (magenta) and Granzyme B (green) lytic granule diameter (nm) for primary CD4 and CD8 T cell. Each dot represents one individual granule. Kruskal-Wallis test with Dunn’s correction, median is shown.

**i, j**, relative frequency (%) distribution of Granzyme B (green) and Perforin (magenta) lytic granules diameter (nm) in primary CD4 (i) and CD8 (j) T cell.

**k**, scheme representing the measurement of polarization index to synaptic dome for Granzyme B and Perforin lytic entities. It is defined as the distance of the granule center of mass to the the synatic dome divided by the distance between the dome and the top of the cell.

**l,m,** average polarization index to dome of all lytic granules in primary CD4 (i) and CD8 (m) T cell plated on non-activating (NA) or activating surface for 5 min. Each dot represents one cell. t test with Welch’s correction, mean± SD are shown.

**n,o**, average proportion of lytic granules below the dome in primary CD4 (n) and CD8 (o) T cell plated on non-activated (NA) or activating surface for 5min. . Each dot represents one cell. t test with Welch’s correction, mean± SEM are shown. ns: not significant, *<0.05, ** <0.01, *** <0.001, **** <0.0001

**Extended Data Figure 5 3D rendering of lytic granules at the IS and tissue expansion**

**a-c** , single z-stack confocal image (a,b) and 3D surface reconstruction (c) of a cryo-fixed and expanded CD8 T cell seeded on activating surface for 5 minutes, and stained for mCLING (grey), Granzyme B (GRZB, blue) and Perforin (PERF, magenta). a, top views (xy axis) showing GRZB and PERF (left) and GRZB, PERF with mCLING (right). b, side view (xz axis) showing GRZB, PERF with mCLING. c, 3D views of the synaptic dome internal surface segmented based on mCLING (transparent grey) signal, PERF (magenta) and GRZB (blue) lytic granules. Orange arrowheads in a,b and c point at the same PERF foci, scale: 2µm.

**d**, 3D surfaces reconstruction of the confocal image presented in figure 5i, of a cryo-fixed and expanded TCR engineered (anti-survivin) human T cell (Tc) and BV173 target cell (Ta) pair labelled for NHS (grey), LAMP1 (orange), Granzyme B (magenta) and Perforin (cyan). The white arrowhead points at GRZB and LAMP1 positive lytic vesicle potentially opening to the synaptic cleft.

**e**, Schematic representation of the workflow for expanding FFPE glioblastoma sections. Non-expanded tissue sections are immunofluorescently labeled for CD4 or CD8 maker and a tile scan of the entire tissue is performed using widefield fluorescent microscope (10x objective). The tissue is then screeded with a 63x objective for regions of interest. A picture is subsequently taken using a 10x objective to map the regions of interest before performing U-ExM procedure on the tissue sample. Gel punches are then made to isolate the previously identified region of interest and stained for CD4, CD8 and protein of interest. After imaging, an additional stripping step can be added for multiplexing.

**f,** expanded human glioblastoma tissue from FFPE sections sequentially multiplexed immunofluorescent labelled and stained for LAMP1 (grey), CD8 (green), CD3 (orange), Perforin (PERF, magenta) and nucleus (DAPI, blue). Microvilli (white arrowhead) are detected as well as the ultrastructure of lytic granules, scale bar: 2 µm.

## Video legends

**Extended Data Video 1**

3D reconstruction of a widefield fluorescent image (Extended Data Fig. 1f) from a cryo-fixed and expanded TCR engineered (anti-survivin) human T cells and BV173 target cell pair stained for NHS (grey) and TOM20 (green) and image with a widefield fluorescent microscope. The movie shows the cell surface (transparent grey) and mitochondria surfaces (magenta) segmentation using NHS signal, the latest correlating with the mitochondria segmentation based on TOM20 (transparent green) signal.

**Extended Data Video 2**

3D reconstruction and surface segmentation of a widefield fluorescent image from a cryo-fixed and expanded primary CD8 T cell plated on activating surface for 5min, stained with NHS (grey), LAMP1 (orange), Granzyme B (cyan) and Perforin (magenta).

**Extended Data Video 3**

3D reconstruction and surface segmentation of one single-core and on multicore granule from a cryo-fixed and expanded CD8 T cell labeled for LAMP1 (white), Granzyme B (GrzB, green), Perforin (PERF, magenta). Wide field fluorescent image (Figure 4e)

**Extended Data Video 4**

3D reconstruction and surface segmentation of a confocal image (figure 4j) from a cryo-fixed and expanded CD8 T cell plated on activating surface for 5min and labeled for mCLING (white), Granzyme B (GrzB, green), Perforin (PERF, magenta).

**Extended Data Video 5**

3D reconstruction of a confocal image from a cryo-fixed and expanded TCR-transgenic human T cells and BV173 target cell pair labeled for NHS (grey), Actine (magenta), tubulin (green) and nucleus (DAPI, cyan).

**Extended Data Video 6**

3D reconstruction of a confocal image (figure 5i) from a cryo-fixed and expanded TCR-transgenic human T cells and BV173 target cell pair labeled for NHS (grey), LAMP1 (orange), Granzyme B (cyan) and Perforin (magenta). This movie focuses on the synaptic cleft.

**Extended Data Video 7**

3D surface segmentation of a confocal image (figure 5i) from a cryo-fixed and expanded TCR-transgenic human T cells and BV173 target cell pair labeled for NHS (transparent and solid grey), LAMP1 (orange), Granzyme B (cyan) and Perforin (magenta). This movie focuses on the synaptic cleft.

**Extended Data Video 8**

3D reconstruction and surface segmentation of a confocal image (supplemental figure 5a) from a cryo-fixed and expanded primary CD8 T cell plated on activating surface for 5min and labeled for mCLING (grey), Granzyme B (cyan) and Perforin (magenta). This movie shows the internal surface of the synaptic dome.

**Extended Data Video 9**

3D reconstruction and surface segmentation of a confocal image (figure 5m) from an expanded FFPE glioblastoma section labeled for CD8 (green), LAMP1 (orange) and Perforin (magenta) and nucleus (DAPI, cyan).

