## Supplementary figures 1 - 5 for "Unveiling the Molecular Architecture of T Cells and Immune Synapses with Cryo-Expansion Microscopy"

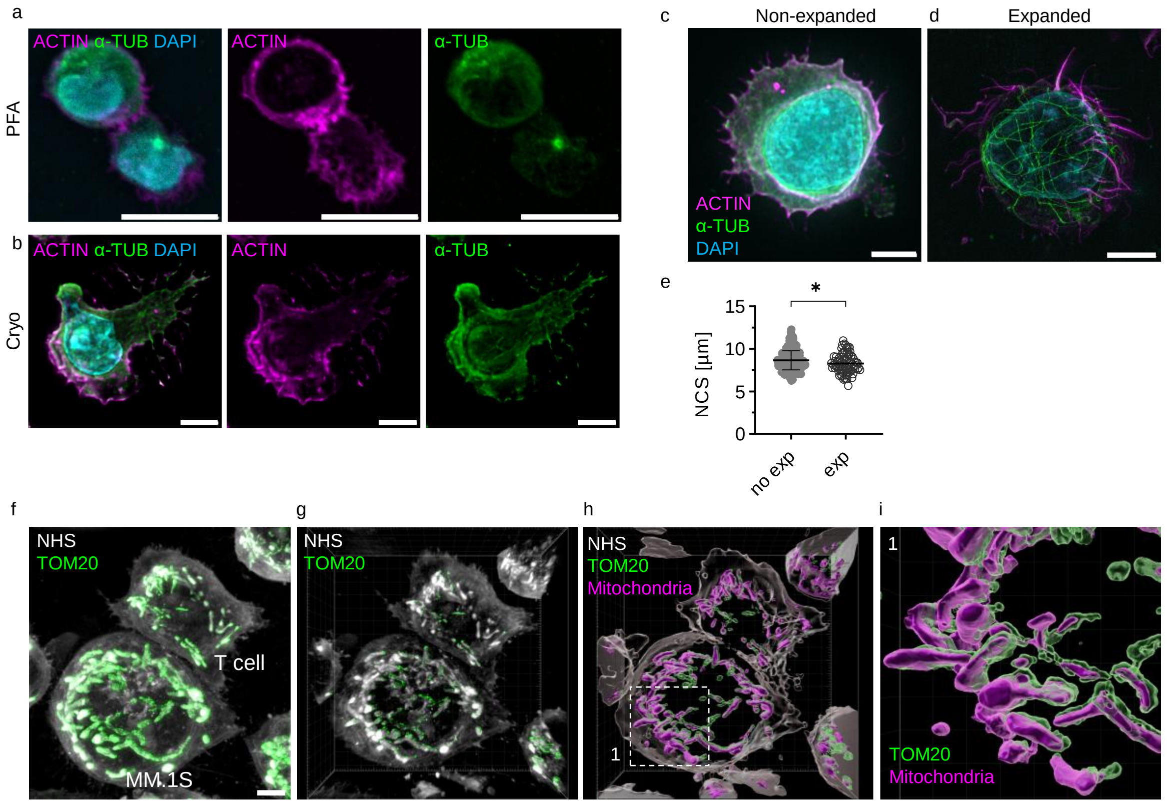


Extended Data Figure 1

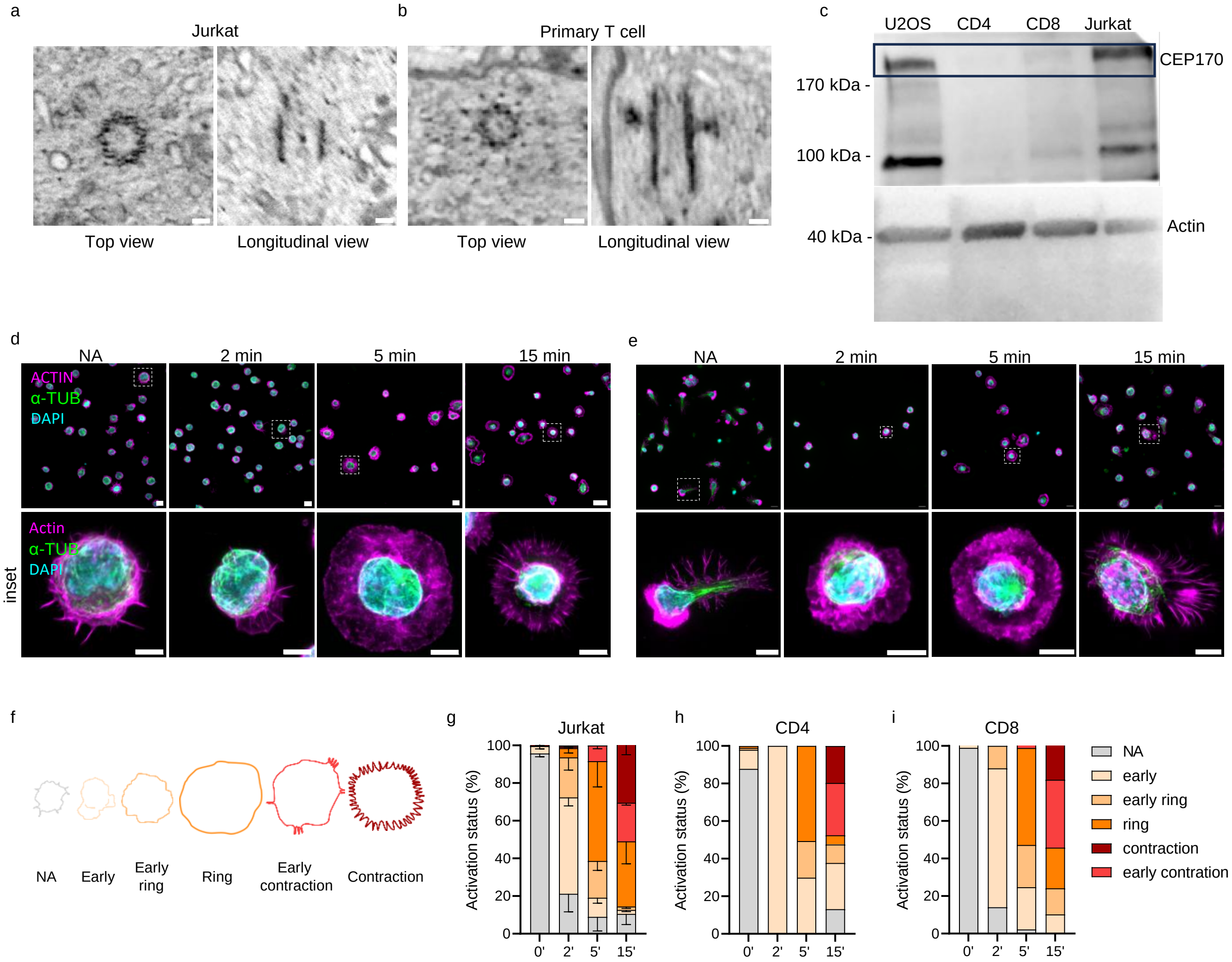


Extended Data Figure 2

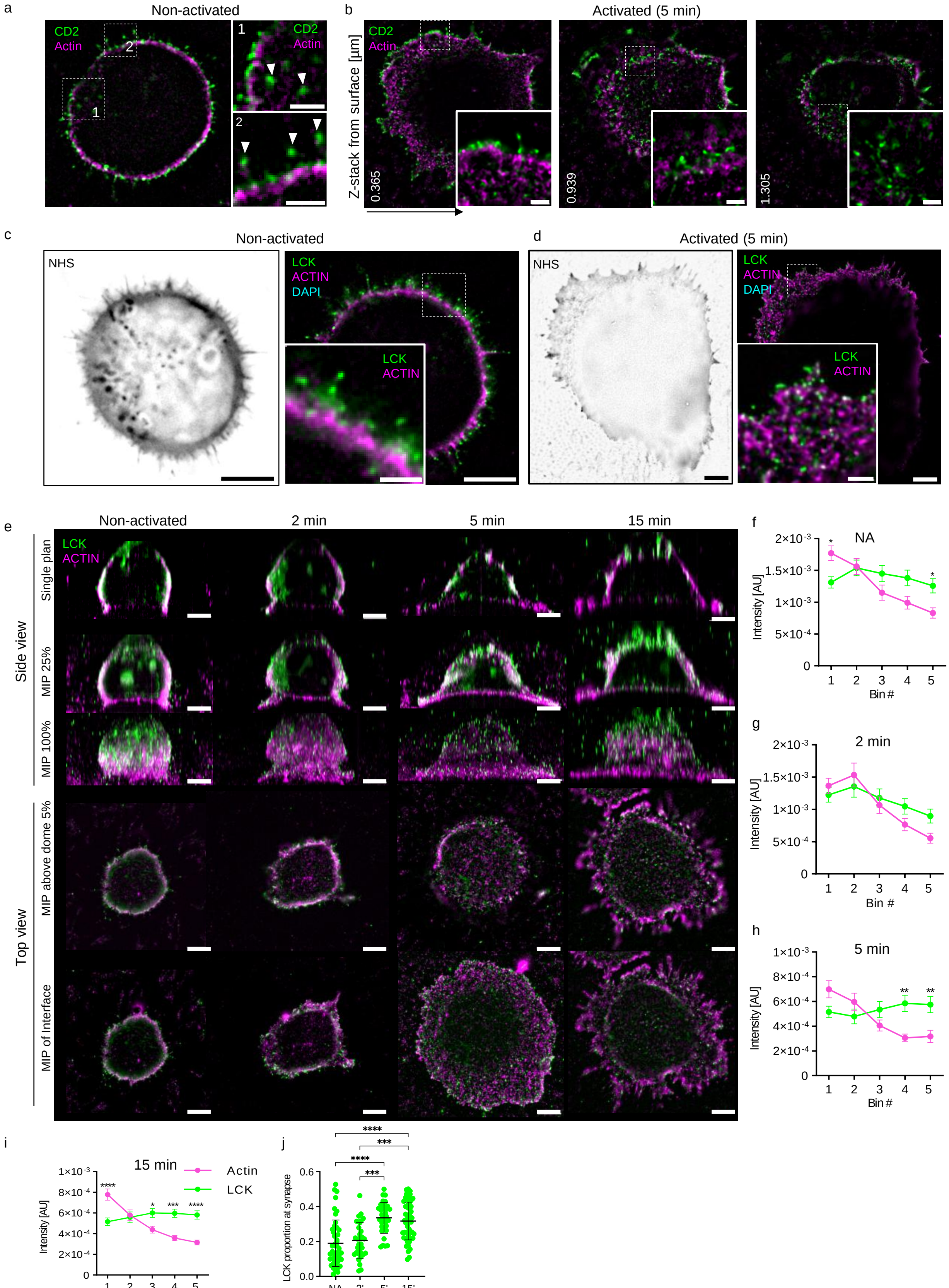


Extended Data Figure 3

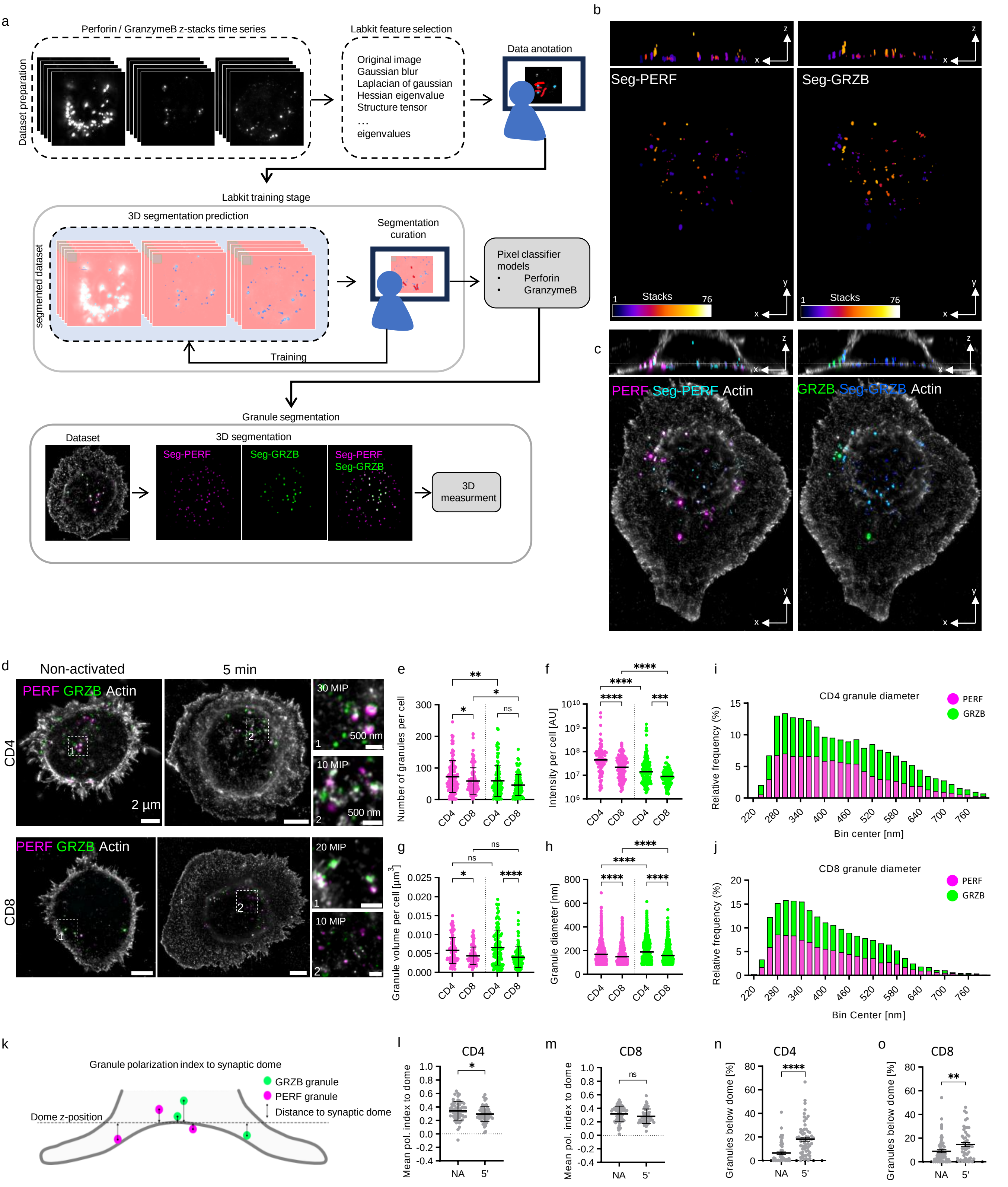


Extended Data Figure 4

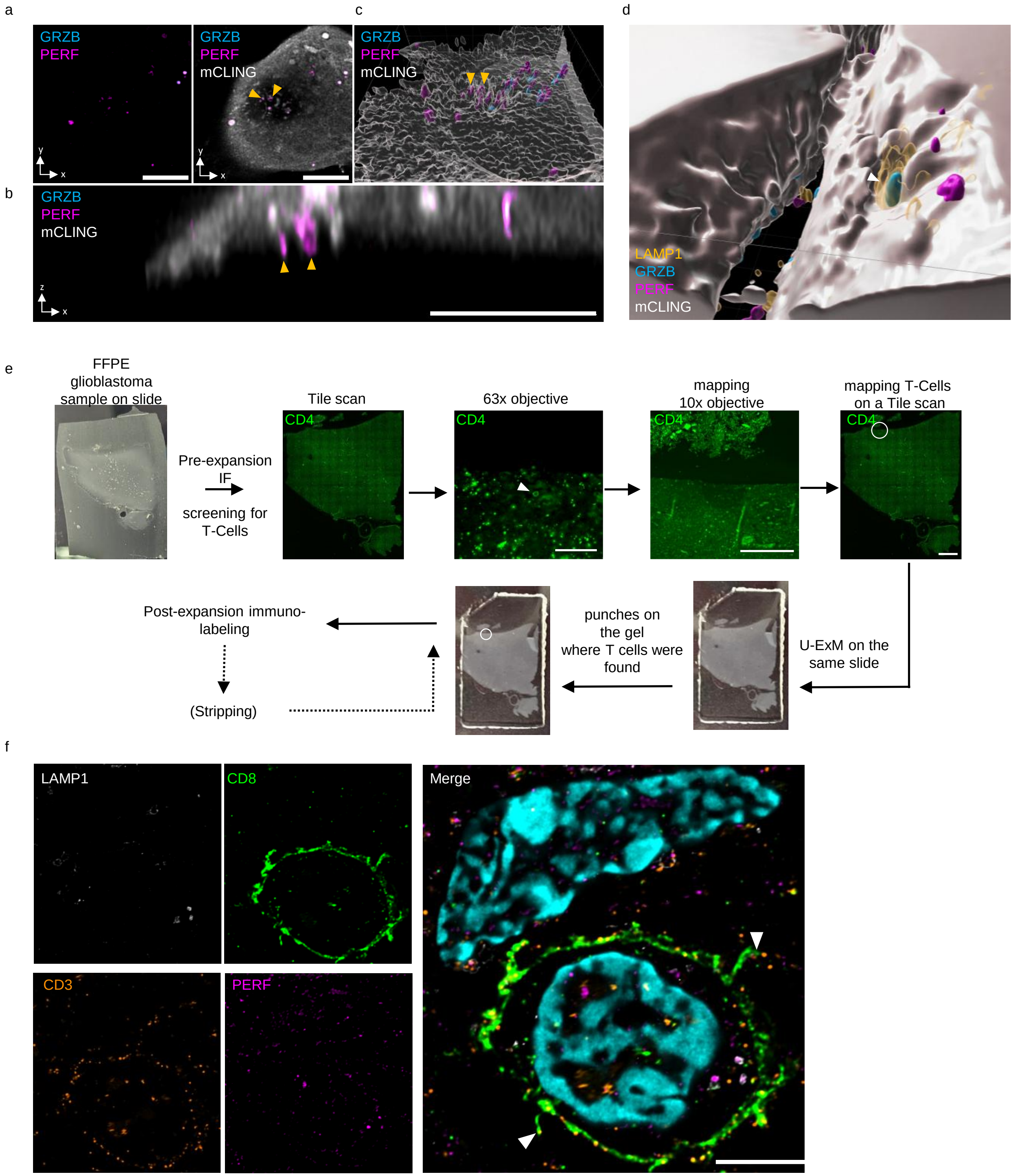


Extended Data Figure 5
